# α_1A_-adrenoceptor inverse agonists and agonists modulate receptor signalling through a conformational selection mechanism

**DOI:** 10.1101/866475

**Authors:** Feng-Jie Wu, Lisa M. Williams, Alaa Abdul-Ridha, Avanka Gunatilaka, Tasneem M. Vaid, Martina Kocan, Alice R. Whitehead, Michael D.W. Griffin, Ross A.D. Bathgate, Daniel J. Scott, Paul R. Gooley

## Abstract

G-Protein Coupled Receptors (GPCRs) transmit signals across the cell membrane via an allosteric network from the ligand-binding site to the G-protein binding site via a series of conserved microswitches. Crystal structures of GPCRs provide snapshots of inactive and active states, but poorly describe the conformational dynamics of the allosteric network that underlies GPCR activation. Here we analyse the correlation between ligand binding and receptor conformation of the α_1A_-adrenoceptor, known for stimulating smooth muscle contraction in response to binding noradrenaline. NMR of ^13^C^ε^H_3_-methionine labelled α_1A_-adrenoreceptor mutants, each exhibiting differing signalling capacities, revealed how different classes of ligands modulate receptor conformational equilibria. ^13^C^ε^H_3_-methionine residues near the microswitches revealed distinct states that correlated with ligand efficacies, supporting a conformational selection mechanism. We propose that allosteric coupling between the microswitches controls receptor conformation and underlies the mechanism of ligand modulation of GPCR signalling in cells.

## Introduction

G-protein coupled receptors (GPCRs) are integral membrane proteins sharing a common seven-helix transmembrane domain (TMD). Conformational changes to the TMD are required to transmit the extracellular stimuli intracellularly to activate signalling pathways. Over the past 20 years X-ray crystal structures, and more recently cryo-EM structures, have revealed a plethora of structural details on how functionally different ligands interact with GPCRs and the conformational changes they induce. Most structures solved to date are of GPCRs in inactive states, bound to inverse agonists or antagonists (Manglik and Kruse, 2017). A few have been crystalized with agonist alone (Manglik and Kruse, 2017), with resultant structures similar to antagonist bound inactive states. Complexes of GPCRs with active-state-stabilising nanobodies, engineered mini G proteins, Gα C-terminal peptide or heterotrimeric G proteins appear necessary to stabilise agonist bound GPCRs in active states for X-ray and cryo-EM structure determination (Carpenter and Tate, 2017). Using these tools, several active state GPCR structures have been solved (Carpenter and Tate, 2017; Garcia-Nafria and Tate, 2019; Manglik and Kruse, 2017), revealing conserved conformational changes that occur upon receptor activation. These include rearrangements in the ligand binding site and a large outward movement at the cytoplasmic side of transmembrane (TM) helix 6 (TM6) to accommodate G protein binding. While providing a wealth of structural detail of static receptor conformations, these structures generally do not provide insight into GPCR signalling complexities such as basal receptor activity, partial agonism and biased agonism.

To address this shortfall, spectroscopic techniques, supported by molecular dynamic simulations, have given insight into the conformational dynamics that underlie the activity of a few diffusible ligand-activated GPCRs including β_2_ adrenergic receptor (β_2_-AR) (Bokoch et al., 2010; Eddy et al., 2016; Horst et al., 2013; Kofuku et al., 2012; Kofuku et al., 2014; Liu, 2012; Manglik et al., 2015; Nygaard et al., 2013), β_1_ adrenergic receptor (β_1_-AR) (Isogai et al., 2016; Solt et al., 2017), adenosine A_2A_ receptor (A_2A_R) (Clark et al., 2017; Eddy et al., 2018; Ye et al., 2018; Ye et al., 2016), µ opioid receptor (µOR) (Okude et al., 2015; Sounier et al., 2015), leukotriene B4 receptor (BLT2)(Casiraghi et al., 2016), and the M2 muscarinic acetylcholine receptor (M2R) (Xu et al., 2019). By far the most studied receptor in this regard is β_2_-AR, for which ^13^C^ε^H_3_-methionine labelling NMR (Bokoch et al., 2010; Kofuku et al., 2012; Kofuku et al., 2014; Nygaard et al., 2013), ^19^F NMR(Eddy et al., 2016; Horst et al., 2013; Liu, 2012; Manglik et al., 2015) and electron paramagnetic resonance (EPR) (Manglik et al., 2015) have been applied to characterise the conformational signatures of this receptor when bound to various ligands and a G protein mimetic nanobody. These studies reveal that GPCRs are highly dynamic, sampling inactive and active conformational states, and are thought to predominantly function via a conformational selection mechanism (Shimada et al., 2018). Such a mechanism posits that a GPCR constantly samples various inactive and active conformations, all existing in equilibrium. Ligands preferentially bind to particular receptor states, depending on their pharmacological characteristics, thus shifting the conformational equilibrium towards these preferred states and modulating the signalling output of the system. The extracellular orthosteric ligand binding site in adrenoceptors is connected to the intracellular G protein binding site through a series of conserved microswitches (Ahuja and Smith, 2009; Deupi and Standfuss, 2011; Trzaskowski et al., 2012) (Figure 1): a central transmission switch (also called the connector region, CWxP motif or PIF motif (Latorraca et al., 2017)), the NPxxY switch, and the intracellular G protein binding site, characterized by the DRY motif (or switch). How these microswitches coordinate the transmission of the extracellular signal is not clear, but molecular dynamics (MD) simulations and NMR data have led to a mechanistic description of “loose allosteric coupling” (Latorraca et al., 2017).

**Figure 1.**
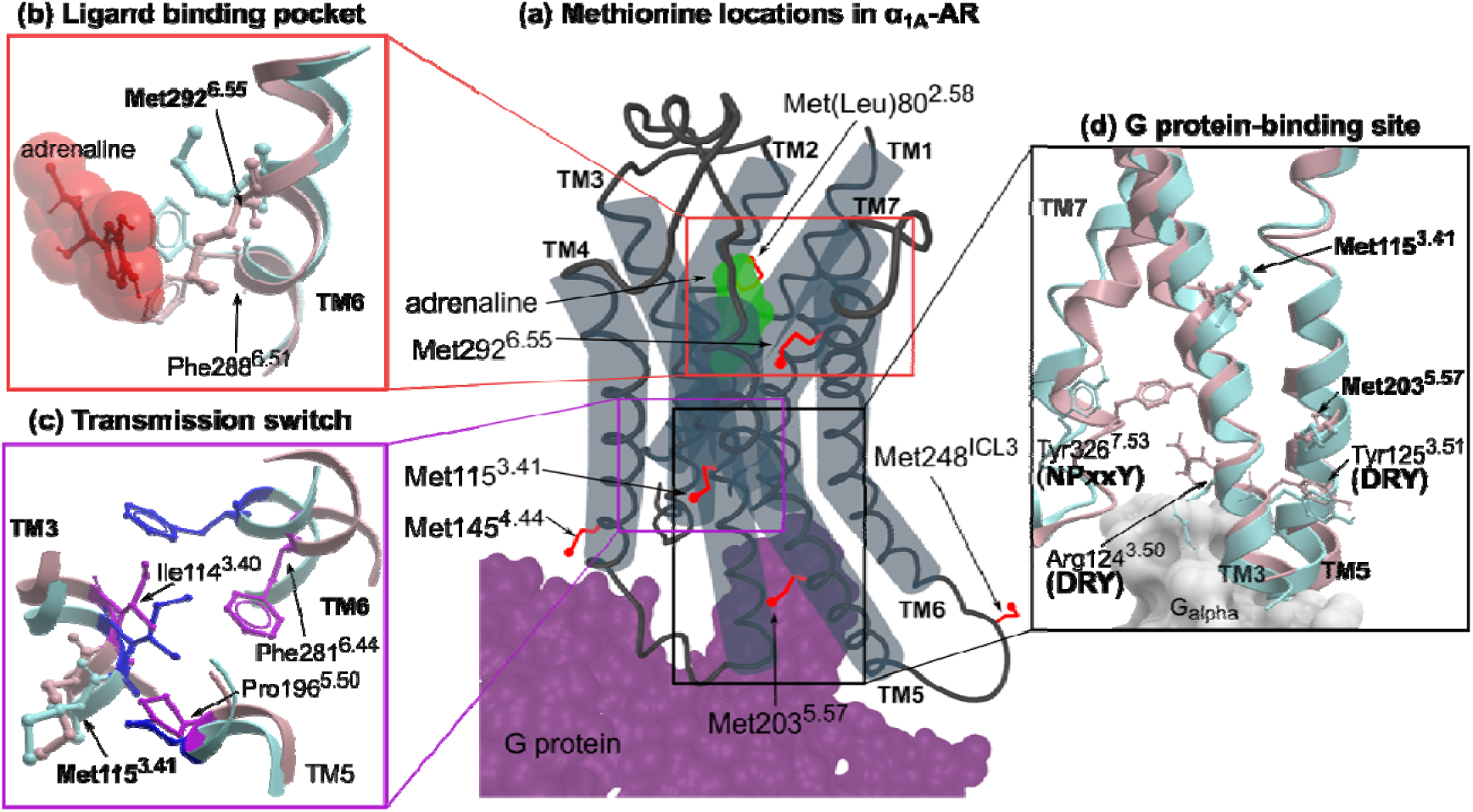
Methionine residues in α_1A_-AR. (a) The location of six methionines on a cartoon representation of α_1A_-AR. Methionine sidechains are highlighted as red sticks. Bound adrenaline and G protein are coloured in green and purple respectively. (b-d) Homology models of α_1A_-AR-A4 in the inactive state (blue; modeled on the X-ray crystal structure of inactive β_2_-AR, pdb id: 5jqh) and active state (pink; modeled on the X-ray crystal structure of active β_2_-AR, pdb id: 3sn6) are superimposed showing inferred conformational changes that occur in the ligand binding pocket (b), transmission switch (c) and G protein binding site (d).

This mechanism refers to each microswitch as conformationally independent from the others, that is an active DRY motif state is not significantly dependent on an active state in the transmission switch. That said, an active state in the transmission switch does increase the probability of the DRY motif (and thus the receptor) to sample active states (thus, loose allosteric coupling) (Latorraca et al., 2017). Put simply, the conformational changes that occur in the microswitches are thought to drive the overall equilibrium state of the receptor system. Despite recent work, it is not well understood how the binding of ligands such as inverse agonists influence the microswitch state equilibria to decrease basal receptor activity.

α_1_-adrenoceptors (α_1_-ARs) comprise three G_q_-coupled GPCR subtypes (α_1A_-, α_1B_- and α_1D_-AR) that bind and sense the endogenous catecholamines, adrenaline and noradrenaline, to modulate a range of physiological processes. In the periphery, postsynaptic α_1_-AR stimulation by catecholamines mediates smooth muscle contraction, thus α_1_-AR antagonists and inverse agonists are clinically prescribed to treat hypertension and benign prostatic hyperplasia (BPH) (Akinaga et al., 2019). α_1_-ARs are also widely expressed in the central nervous system (CNS), but the lack of subtype-selective antibodies and ligands limits the understanding of their role in neuroplasticity and neurodegeneration (Perez and Doze, 2011). Currently there are no available crystal structures of an α_1_-AR family member, which limits the rational design of more selective compounds to probe the physiological role of α_1A_-AR in the CNS.

Recombinant α_1A_-AR expresses poorly and the resultant protein is particularly unstable when purified in detergent (Scott and Pluckthun, 2013), which has hindered biochemical studies of this GPCR. Recently, we engineered an α_1A_-AR variant, α_1A_-AR-A4, that can be expressed in *Escherichia coli* (*E. coli*) and exhibits improved stability when purified in detergents (Yong et al., 2018). When expressed in COS-7 cells α_1A_-AR-A4 exhibits no signalling efficacy in response to adrenaline stimulation (Yong et al., 2018). In the present study, α_1A_-AR-A4 was labelled with ^13^C^ε^H_3_-methionine at the five naturally occurring methionine residues, providing NMR probes to assess how inverse agonists, partial agonists and full agonists influence receptor conformational equilibria. Three of these methionines are excellent probes of the ligand-binding site and the microswitches proposed to be markers of signal transmission: Met292^6.55^ (superscript denotes GPCRdb numbering (Isberg et al., 2015)) is located in the ligand binding site; Met115^3.41^ is proximal to the transmission switch (Ile114^3.40^, Pro196^5.50^, Leu197^5.51^, Phe281^6.44^, Trp285^6.48^); and Met203^5.57^ sits above the tyrosine of the DRY motif (Asp130^3.49^, Arg131^3.50^, Tyr132^3.51^). Using the inactive α_1A_-AR variant, α_1A_-AR-A4, and by reverse mutation to an active receptor (α_1A_-AR-A4-active) we show that for Met115^3.41^ and Met203^5.57^ the chemical shifts and line-widths of the ^13^C^ε^H_3_ groups are dependent on ligand efficacy (from strong inverse agonist to full agonist), suggesting that α_1A_-AR activation proceeds primarily through a conformational selection mechanism.

## Results

### ^13^C^ε^H_3_ methionine labelling and NMR signal assignment

α_1A_-AR-A4 is a thermostabilised variant of the human α_1A_-AR that contains 15 amino acid substitutions over wild type (WT) human α_1A_-AR (Supplementary Figure 1). Excluding Met1, α_1A_-AR-A4 possesses six methionine residues, five of which are naturally occurring (Met115^3.41^, Met145^4.44^, Met203^5.57^, Met248^ICL3^, Met292^6.55^) and one, Met80^2.58^, is a thermostabilising mutation previously selected for (Yong et al., 2018) (Supplementary Figure 1). Homology models of α_1A_-AR (Figure 1) built on in inactive- and active-states of X-ray structures of β_2_-AR show that three of these methionines were particularly interesting as conformational probes as they are located either within the adrenaline binding site (Met292^6.55^), immediately adjacent to the highly conserved Ile114^3.40^ of the transmission switch (Met115^3.41^), or sitting above Tyr125^3.51^ of the DRY motif within the G protein binding site (Met203^5.57^). These homology models of α_1A_-AR suggest that each of these regions undergo significant local rearrangements between inactive to active conformations (Figure 1).

α_1A_-AR-A4 was expressed and labelled with ^13^C^ε^H_3_-methionine using an adapted *E. coli* methionine biosynthesis pathway inhibition protocol that we have previously used to generate ^13^C^ε^H_3_-methionine-labeled neurotensin receptor 1 (NTS_1_) samples labelled with 96% incorporation efficiency(Bumbak et al., 2019; Bumbak et al., 2018). Using this method α_1A_-AR-A4 expressed well and could be purified, solubilized in n-dodecyl β-D-maltopyranoside (DDM), with a yield of (0.5-1 mg/L culture). 40-60 µM samples of ^13^C^ε^H_3_-methionine-labeled α_1A_-AR-A4 were subsequently used to record 2D ^1^H-^13^C SOFAST-heteronuclear multiple quantum coherence (HMQC) spectra in the apo state, and in the bound states for prazosin (full inverse agonist), WB-4101 (partial inverse agonist), phentolamine (partial inverse agonist), silodosin (or KMD-3213, neutral antagonist), oxymetazoline (partial agonist) and adrenaline (full agonist) (Figure 2 and Supplementary Figure 2). Individual ^13^C^ε^H_3_-methionine resonances were assigned by expressing and analysing α_1A_-AR-A4 M80L, α_1A_-AR-A4 M115I, α_1A_-AR-A4 M203L, α_1A_-AR-A4 M248I and α_1A_-AR-A4 M292I mutants in the same way. ^1^H-^13^C SOFAST-HMQC spectra enabled clear assignment of mutated methionines as the remaining five resonances in these spectra showed only small chemical shift differences in the presence of the mutation (Supplementary Figure 3). The ^13^C^ε^H_3_-methionine of the apo state of α_1A_-AR-A4 showed clear single resonances for each methyl with no significant heterogeneity, in contrast to many previously studied GPCRs (Casiraghi et al., 2016; Kofuku et al., 2012; Nygaard et al., 2013; Okude et al., 2015; Solt et al., 2017; Xu et al., 2019), (Figure 2). Met145^4.44^ and Met248^ICL3^ exhibited intense signals with ^1^H and ^13^C chemical shifts of the methyl group indicative of solvent exposed, unrestrained methyl groups. Met248^ICL3^, located within ICL3 (Figure 1a), showed strong signal intensity most likely due to the mobility of this loop and exposure to the bulk solvent. Met145^4.44^ is at the C-terminal intracellular end of TM4, predicted to be exposed on the surface of the helix (Figure 1a) and thus also highly mobile. Met80^2.58^ was not unambiguously assigned (Supplementary Figure 3a,f) as it either is significantly broadened and difficult to resolve in all receptor states or may overlap with Met145^4.44^ and under some conditions with Met292^6.55^ (Supplementary Figure 3e). The remaining methionines, Met115^3.41^, Met203^5.57^ and Met292^6.55^, were readily assigned (Supplementary Figure 3b,c,e,g,h,j) and exhibited resolved chemical shifts for the ^13^C^ε^H_3_ that were sensitive to the bound ligand (Figure 2b-d).

**Figure 2.**
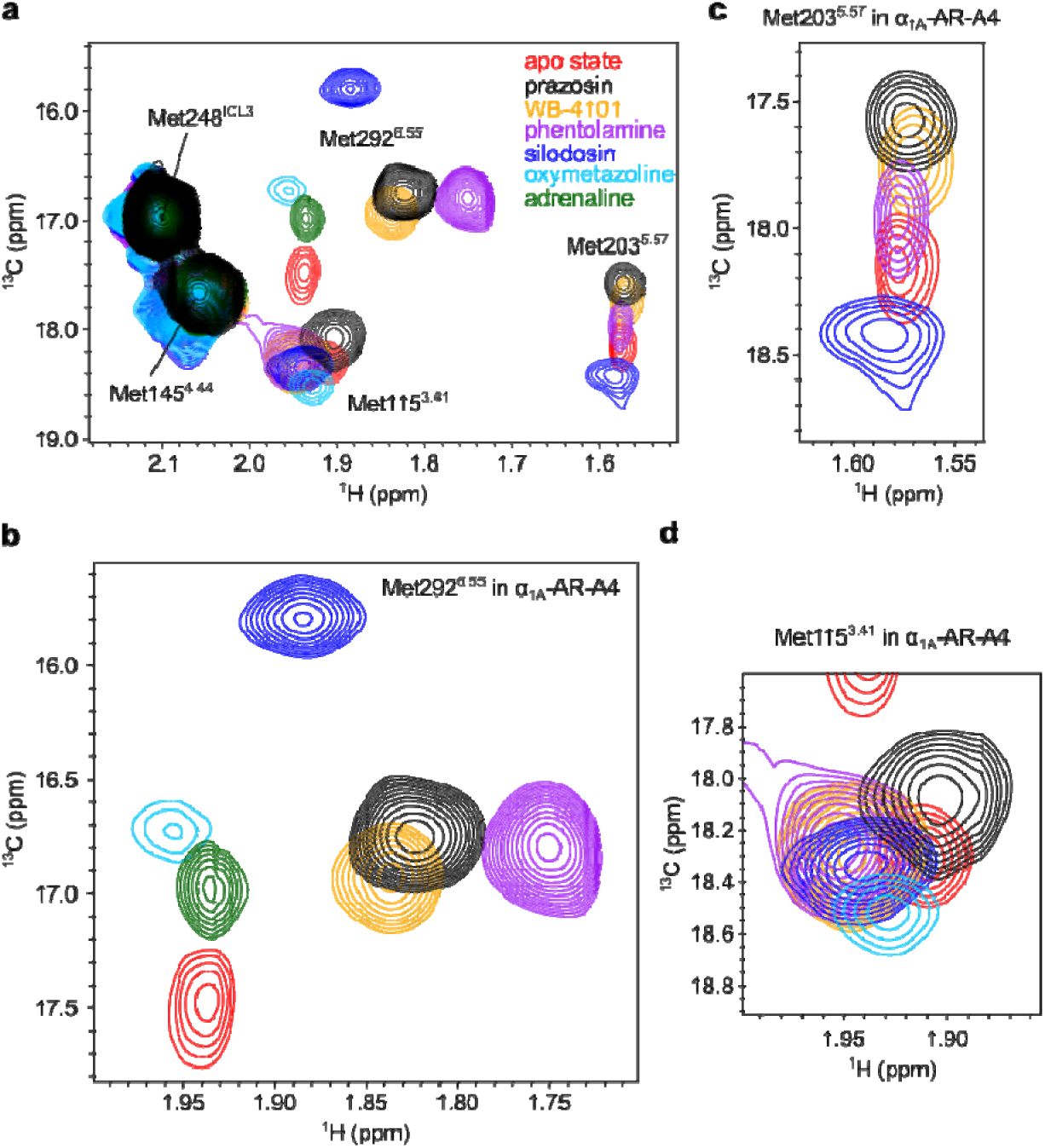
^1^H-^13^C SOFAST-HMQC spectra of α_1A_-AR-A4. (a) Overlay of 2D ^1^H-^13^C SOFAST-HMQC spectra for [^13^C^ε^H_3_-Met] α_1A_-AR-A4 collected in the apo state (red) and bound to prazosin (black, inverse agonist), WB-4101 (yellow, inverse agonist), phentolamine (purple, inverse agonist), silodosin (blue, neutral antagonist), oxymetazoline (cyan, partial agonist) and adrenaline (green, full agonist). (b) Close-up of the Met292^6.55^ resonance. (c) Close-up of the Met203^5.57^ resonance. (d) Close-up of the Met115^3.41^ resonance. Spectra were acquired on ∼50 µM α_1A_-AR-A4 dissolved in 0.02-0.1% DDM micelle, pH 7.5 and 25 °C.

Based on homology models, Met292^6.55^ projects into the orthosteric ligand binding pocket (Figure 1a) and mutational studies support a role for this residue in ligand binding (Hwa et al., 1995). Thus, the ^13^C^ε^H_3_ chemical shifts of Met292^6.55^ likely reflect a direct interaction with chemical groups of each ligand. Interestingly, the resonance intensities of Met292^6.55^ increased in the presence of antagonists and inverse agonists relative to the apo state (Figure 2b), indicating that binding of these ligands reduces conformational dynamics in the orthosteric binding site. Met115^3.41^ and Met203^5.57^ are distant from the orthosteric site, but both the chemical shifts and linewidths of their ^13^C^ε^H_3_ groups were sensitive to ligand binding (Figure 2c,d), likely reflecting receptor conformational changes in the transmission switch and G protein-binding site respectively (Figure 1c,d). The ^1^H chemical shift of the methyl of Met203^5.57^ was shifted upfield from typical small-peptide positions (2.1 ppm) to 1.58 ppm in agreement with our models, which predict ring-current induced effects from Tyr125^3.51^ of the DRY motif (Supplementary Figure 4). Met203^5.57^ therefore serves as a probe of conformational change within this region. Indeed, the resonances of the ^13^C^ε^H_3_ of Met203^5.57^ exhibited a significant linear chemical shift change depending on which ligand was bound, demonstrating that allosteric coupling between the ligand binding site and the G protein-binding site is retained in the inactive α_1A_-AR-A4 in solution. Such a linear chemical shift change is also expected for ligands modulating receptor state *via* conformational selection. We postulate that the Met203^5.57^ signal reflects the average, equilibrium signal, between inactive and active states undergoing fast exchange. Inverse agonists preferentially bound to inactive states, shifting the Met203^5.57^ equilibrium to an upfield position (inactive state) compared to apo state receptor, which can sample active-like states to a certain degree.

We were interested to see if an opposite trend could be observed for receptor agonists, which we hypothesised would shift the position of the Met203^5.57^ resonance downfield. For α_1A_-AR-A4 however, the binding of the full agonist adrenaline to α_1A_-AR-A4 resulted in complete line broadening of the Met115^3.41^ and Met203^5.57^ resonances despite the promotion of a distinct chemical shift for Met292^6.55^ in the binding site. Binding of the partial agonist oxymetazoline resulted in substantial broadening of Met203^5.57^ and Met292^6.55^, but not Met115^3.41^. The loss of these chemical shifts upon agonist binding was likely due to the significantly weaker agonist affinities at α_1A_-AR-A4 compared to unmutated, WT α_1A_-AR, as a result of the F312L stabilizing mutation (Yong et al., 2018). Thus, NMR experiments were repeated on α_1A_-AR-A4 (L312F), for which agonist affinities were largely restored to that of WT α_1A_-AR (Supplementary Figure 5 and Supplementary Table 1) (Yong et al., 2018).

### Agonist induced chemical shifts of Met115^3.41^ and Met203^5.57^ resonances

Despite the reduced thermostability of α_1A_-AR-A4 (L312F) (Yong et al., 2018), we were able to ^13^C^ε^H_3_-methionine-label and record ^1^H-^13^C SOFAST-HMQC spectra for this receptor in the apo state and bound to adrenaline (full agonist), phenylephrine (full agonist), A-61603 (full agonist), and oxymetazoline (partial agonist) in addition to the inverse agonists and neutral antagonists tested on α_1A_-AR-A4 (Figure 3a). Overall the ^1^H-^13^C SOFAST-HMQC spectra of the apo, antagonist and inverse agonist bound states of α_1A_-AR-A4 (L312F) were similar to those of α_1A_-AR-A4. Again, single resonances for the ^13^C^ε^H_3_-methionine groups of α_1A_-AR-A4 (L312F) were observed for all ligands. The chemical shifts of Met292^6.55^ induced by each ligand in α_1A_-AR-A4 (L312F) were slightly different to those of α_1A_-AR-A4, most likely due to orthosteric binding site changes after the L312F reversion. Inverse agonist binding increased the intensity of the Met292^6.55^ resonance in α_1A_-AR-A4 (L312F), as was seen with α_1A_-AR-A4; whereas the neutral antagonist silodosin significantly decreased the peak intensity and the partial agonist oxymetazoline and full agonist A-61603 highly broadened the resonance of Met292^6.55^ in α_1A_-AR-A4 (L312F) (Supplementary Figure 6).

**Figure 3.**
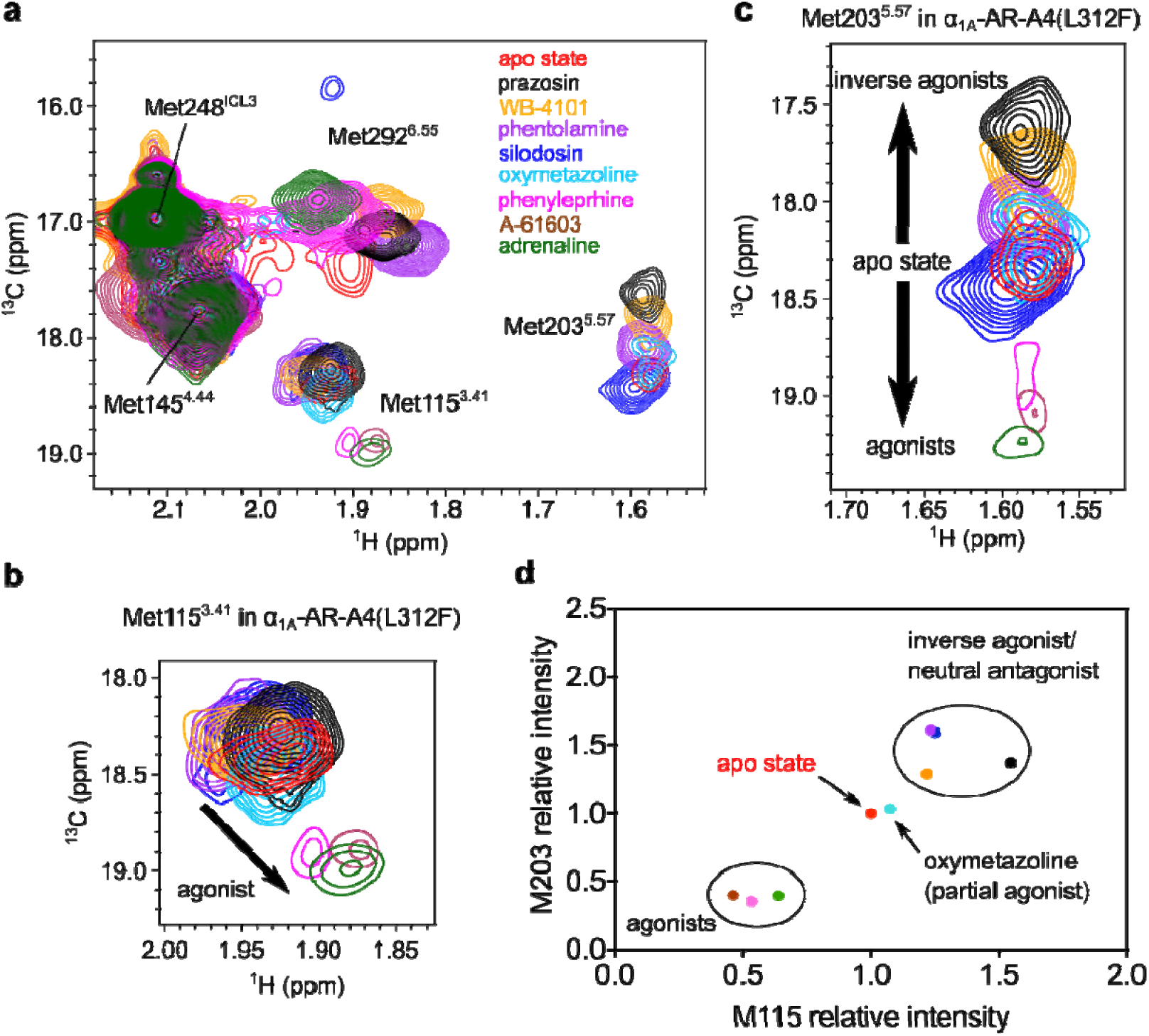
Ligand efficacy-dependent chemical shifts of Met115^3.41^ and Met203^5.57^ resonances. (a) Overlay of 2D ^1^H-^13^C SOFAST-HMQC spectra for [^13^C^ε^H_3_-Met] α_1A_-AR-A4 (L312F) in the apo state (red) and bound to ligands: prazosin (black, inverse agonist), WB-4101 (orange, inverse agonist), phentolamine (purple, inverse agonist), silodosin (blue, neutral antagonist), oxymetazoline (cyan, partial agonist), phenylephrine (magenta, full agonist), A-61603 (maroon, full agonist), adrenaline (green, full agonist). (b) Close-up of the Met115^3.41^ resonance in α_1A_-AR-A4 (L312F). (c) Close-up of the Met203^5.57^ resonance in α_1A_-AR-A4 (L312F). The spectra for adrenaline, A-61603 and phenylephrine are plotted at a level 1.8-times lower than the main figure. (d) Normalized peak intensities of Met115^3.41^ and Met203^5.57^ of α_1A_-AR-A4 (L312F) show differences between agonists, antagonists and partial agonists. Ligands are coloured as listed above. Spectra were acquired on ∼50 µM α_1A_-AR-A4 (L312F) dissolved in 0.02-0.1% DDM micelle, pH 7.5 and 25 °C.

The recovered agonist affinity for α_1A_-AR-A4 (L312F) allowed the measurement of ^1^H-^13^C SOFAST-HMQC spectra where we were confident of full receptor-agonist saturation. Binding of the full agonist adrenaline to α_1A_-AR-A4 (L312F) produced a similar Met292^6.55^ chemical shift to that seen with α_1A_-AR-A4 (Figure 3a), and also weak peaks were now observed for Met115^3.41^ and Met203^5.57^ (Figure 3b,c), which were completely broadened in adrenaline-bound α_1A_-AR-A4. Importantly, the binding of all agonists, adrenaline, phenylephrine and A-61603 to α_1A_-AR-A4 (L312F) induced distinct chemical shift and line broadening changes to Met115^3.41^ and Met203^5.57^ compared to neutral antagonists and inverse agonists (Figure 3b,c). The agonist-induced Met115^3.41^ resonances cluster together potentially indicative of an active transmission switch conformation (Figure 3b). Binding of the partial agonist oxymetazoline induced a chemical shift of Met115^3.41^ falling between the inverse agonist and full agonist clusters, consistent with partial agonists promoting a weaker shift in the inactive-active transmission switch state equilibrium. The linear change in Met203^5.57^ chemical shift position upon inverse agonist binding seen with α_1A_-AR-A4 was retained in α_1A_-AR-A4 (L312F), but as hypothesised, agonist binding promoted opposite, downfield resonance shifts in the ^13^C dimension along the same vector (Figure 3c). This change in ^13^C^ε^H_3_-methionine chemical shift in the ^13^C dimension reflects a change in the χ3 dihedral angle. The ^13^C chemical shift dependence of this angle is about 19 ppm for trans and 16 ppm for ±gauche (Butterfoss et al., 2010). For the apo and antagonist states the chemical shift of 18.25 to 18.5 ppm suggests a trend toward trans, whereas in the full-inverse agonist state the resonance shifts up-field to 17.7 ppm, indicative of an averaging between gauche and trans. Consistent with our homology models (Supplementary Figure 4) for full agonist a further downfield shift between 19 to 19.25 ppm infers an increase in the trans conformer. Interestingly, the partial agonist oxymetazoline induced a small upfield ^13^C^ε^ shift of Met203^5.57^, similar to the inverse agonist phentolamine. The fact that full agonists induced Met203^5.57^ chemical shifts to move in the opposite direction to inverse agonists suggests an equilibrium shift away from inactive to active conformational states of the DRY motif. Furthermore, the resonance intensities of both Met115^3.41^ and Met203^5.57^ in α_1A_-AR-A4 (L312F), relative to the ligand-insensitive Met145^4.44^ resonance, were weakened upon agonist binding compared to the intensity increases seen with antagonists and inverse agonists (Figure 3d). The intensities of Met115^3.41^ and Met203^5.57^ upon binding of the partial agonist oxymetazoline fell in between the antagonist- and agonist-induced intensities. The behaviour of the ^13^C^ε^H_3_ of Met115^3.41^ and Met203^5.57^ is consistent with the current concept that agonists increase conformational heterogeneity in GPCRs, where agonists increase microsecond timescale transitions to active receptor states, to increase the probability of engaging and activating effector proteins (Kofuku et al., 2012; Manglik et al., 2015; Nygaard et al., 2013; Shimada et al., 2018; Solt et al., 2017; Ye et al., 2016).

### Chemical shift changes of Met203^5.57^ correlate with ligand efficacy

In mammalian cells, α_1A_-AR exhibits basal activity in the absence of bound ligands(Zhu et al., 2000). Such basal activity is unaffected by the binding of neutral antagonists but is reduced by the binding of inverse agonists to the receptor. In the case of α_1A_-AR, by probing the ability of various antagonists to reduce the signalling of a constitutively active receptor mutant, the rank order of inverse agonist efficacies has been found to be: prazosin (strongest); WB-4101; phentolamine (weakest); and silodosin being a neutral antagonist (Zhu et al., 2000). To understand how the NMR signals of Met203^5.57^ in α_1A_-AR-A4 (L312F) relate to receptor conformational equilibria, the changes to the chemical shifts for the ^13^C^ε^H_3_ of Met203^5.57^ were plotted against the previously published relative efficacy values for the inverse agonists, revealing a strong linear correlation (R^2^ = 0.99, Figure 4a). To test if this correlation is retained when probing inverse agonism at the wild-type α_1A_-AR, we determined the relative inverse agonist efficacies of these ligands using a NanoBiT split luciferase assay (Inoue et al., 2019). In this assay the 18 kDa Large BiT (LgBiT) fragment was fused to the N-terminus of Gα_q_ and the 1.3 kDa Small BiT (SmBiT) was fused to the N-terminus of Gγ_2_. When co-expressed with Gβ_1,_ the formation of a Gα_q_(LgBiT)-Gβ_1_-Gγ_2_(SmBiT) heterotrimer results in bright luminescence. GPCR-induced stimulation of this G protein complex causes dissociation of the heterotrimer and thus reduction in luminescence output, whereas inhibition of basal GPCR activation would be predicted to increase luminescence output. COS-7 African green monkey kidney cells stably expressing wild-type (WT) α_1A_-AR were transfected with Gα_q_(LgBiT), Gβ_1_ and Gγ_2_(SmBiT) encoding expression plasmids, incubated with luminescence substrate, and then treated with various α_1A_-AR ligands while monitoring cellular luminescence. A-61603 induced α_1A_-AR activation led to heterotrimer dissociation of the Gα_q_(LgBiT)-Gβ_1_-Gγ_2_(SmBiT) complex and thus a reduction in luminescence output (Figure 4b). Inverse agonists on-the-other-hand reduced basal activation of α_1A_-AR, maintaining the Gα_q_(LgBiT)-Gβ_1_-Gγ_2_(SmBiT) complex leading to increase luminescence output from the cells (Figure 4b). The specificity of these responses was probed by conducting the same experiments on COS-7 cells not expressing α_1A_-AR (Supplementary Figure 7a-c). The observed changes in luminescence after α_1_-AR ligand treatments were specific to α_1A_-AR expressing cells except for WB-4101, which induced a short (5 min) increase in luminescence in the control cells (Supplementary Figure 7a). To exclude this non-specific effect the net luminescence change for each sample group was calculated as the area under the luminescence curves between 5 and 10 min after ligand addition. A strong linear correlation was found between the chemical shift changes for the ^13^C^ε^H_3_ of Met203^5.57^ in α_1A_-AR-A4 (L312F) and the net luminescence increase generated by each inverse agonist over the five-minute period in the WT α_1A_-AR-expressing cells (R^2^ = 0.72, Figure 4c). Importantly, the Met203^5.57^ chemical shift positions of α_1A_-AR-A4 (L312F) did not correlate with the affinity of these antagonists for α_1A_-AR (Figure 4d), demonstrating that the differences in chemical shift were not due to varying receptor occupancy. Furthermore, no correlation was seen between the ^13^C^ε^H_3_ Met203^5.57^ chemical shift changes of α_1A_-AR-A4 (L312F) and the net luminescence changes in COS-7 cells not expressing WT α_1A_-AR (Supplementary Figure 7b). Critically, the correlation between chemical shift changes of ^13^C^ε^H_3_ Met203^5.57^ in α_1A_-AR-A4 (L312F) and WT α_1A_-AR-specific luminescence increases in the NanoBiT assay remained when the analysis window was extended to include the full 10 minutes after ligand addition (Supplementary Figure 7d).

**Figure 4.**
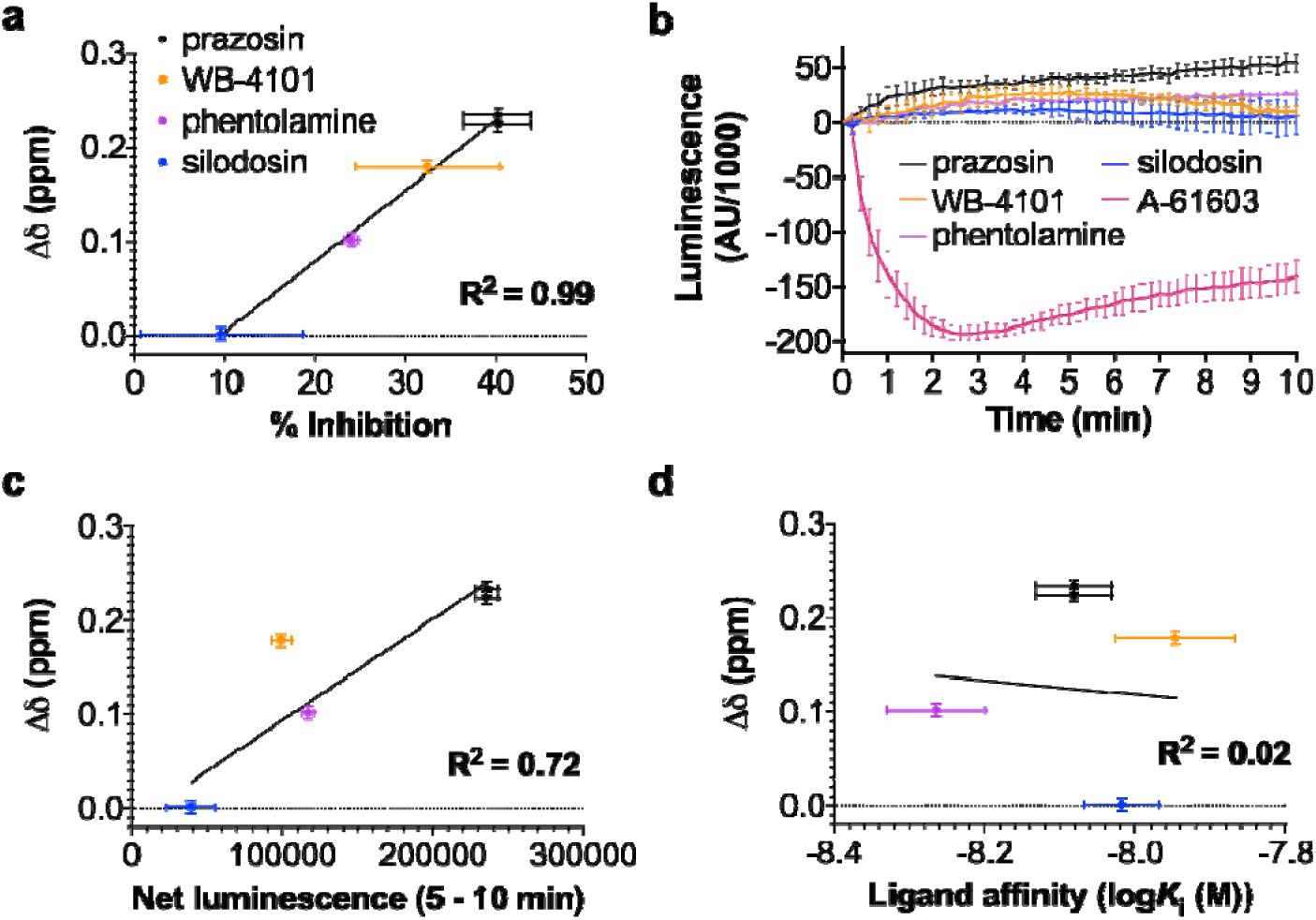
Correlation between the chemical shift positions of the ^13^C^ε^H_3_ in Met203^5.57^ and inverse agonists efficacy. (a) Linear regression analysis of the average chemical shift differences (Δδ) for the ^13^C^ε^H_3_ of Met203^5.57^ in α_1A_-AR-A4 (L312F) when bound to prazosin (black circles), WB-4101 (orange circles), and phentolamine (purple circles) compared to silodosin (blue circles) and the published efficacy of each ligand in reducing the signaling of a constitutively active mutant of α_1A_-AR (Zhu et al., 2000). Published data were extracted using WebPlotDigitizer (https://automeris.io/WebPlotDigitizer). Testing the resultant equation against the null hypothesis of a slope of zero resulted in a P value of < 0.0001 (b) NanoBit G protein activity assay demonstrating inverse agonism of prazosin, WB-4101, phentolamine and silodosin at WT α_1A_-AR-expressing COS-7 cells. Each of these inverse agonist experiments were repeated in three independent biological replicate experiments, with the mean ± SEM of the resultant luminescence plotted for each timepoint. To demonstrate the response from an agonist, A-61603 treatment was performed in two independent biological replicate experiments. Each biological replicate comprised three technical replicates measured in parallel. The grey shaded region indicates where the area under each biological replicate curve was calculated for (c). (c) Linear regression analysis of the average chemical shift differences (Δδ) for the ^13^C^ε^H_3_ of Met203^5.57^ in α_1A_-AR-A4 (L312F) and the increase in luminescence seen in the NanoBit assay for each inverse agonist and neutral antagonist. Ligands are coloured as listed above and the P value testing against a slope of 0 was 0.011 (d) Linear regression analysis of the average chemical shift differences (Δδ) for the ^13^C^ε^H_3_ of Met203^5.57^ in α_1A_-AR-A4 (L312F) and the affinities of each inverse agonist and neutral antagonist. Ligands are coloured as listed above and the P value testing against a slope of 0 was 0.89. In (a), (c) and (d) Δδ are plotted for two independent titrations of prazosin and silodosin, and single experiments for WB-4101 and phentolamine. Average chemical shift differences (Δδ) were normalised using the equation Δδ=[(Δδ_1H_)^2^+(Δδ_13C_/3.5)^2^]^0.5^ and error were calculated by the formula [Δδ_1H_*R_1H_+Δδ_13C_*R_13C_/(3.5)^2^]/Δδ, where R_1H_ and R_13C_ are the digital resolutions in ppm in the ^1^H and ^13^C dimensions respectively (Kofuku et al., 2012).

### Improving signalling competency in α_1A_-AR-A4

When expressed in COS-7 cells, α_1A_-AR-A4 is incapable of stimulating cellular increases in IP_1_ in response to adrenaline binding, or activation of a cyclic adenosine monophosphate (cAMP) response element (CRE) reporter gene after treatment with another α_1_-AR agonist, phenylephrine (Yong et al., 2018). To ensure biological relevance of our NMR studies we thus sought α_1A_-AR-A4 back-mutants that were able to stimulate canonical signalling pathways in mammalian cells upon agonist treatment. Seven thermostabilising mutations within the TMD of α_1A_-AR-A4 were back-mutated (Y67N, M80L, A127G, F151W, K322N, L327P and Y329S) as single changes or in combinations, and screened for phenylephrine and oxymetazoline induced signalling with an IP_1_ assay in transfected COS-7 cells (Supplementary Figure 8a). While the back-mutant, α_1A_-AR-A4 (Y67N, M80L, K322N, L327P, Y329S) was able to facilitate significant oxymetazoline-induced cellular accumulation of IP_1_ compared to α_1A_-AR-A4 and WT α_1A_-AR (Supplementary Figure 8a) it expressed poorly in bacteria. The back-mutant α_1A_-AR-A4 (Y67N, K322N), termed α_1A_-AR-A4-active, however was able to stimulate IP_1_ accumulation in response to both phenylephrine and oxymetazoline treatment (Supplementary Figure 8a) and it expressed well in bacteria. Importantly, α_1A_-AR-A4 contains the N322K-stabilising mutation in the NPxxY switch(Trzaskowski et al., 2012), which is hypothesised to form a stabilizing salt bridge with Asp72^2.50^ to lock the NPxxY switch in an inactive and stable state. We thus expected that reversion of this mutation (K322N) would restore the function of the NPxxY switch and the signalling activity of α_1A_-AR-A4. Interestingly, the Y67N mutation was required on top of K322N to restore signalling activity in α_1A_-AR-A4-active. N67^2.45^ is distant from the NPxxY switch and its importance is not clear.

Using an intracellular calcium mobilisation assay, α_1A_-AR-A4-active was able to be activated by the full agonists adrenaline and A-61603, as well as the partial agonists oxymetazoline and PF-3774076 (Supplementary Figure 8b-e). The affinity of QAPB for α_1A_-AR-A4-active was retained upon purification of the receptor in DDM (Supplementary Figure 9a). Competition binding assays revealed however, that the affinities of agonists for α_1A_-AR-A4-active (Supplementary Figure 9b and Supplementary Table 1) were weaker than WT α_1A_-AR due to the F312L stabilizing mutation, but stronger than at α_1A_-AR-A4 (Yong et al., 2018). When purified in DDM α_1A_-AR-A4-active was significantly less stable than α_1A_-AR-A4 (Supplementary Figure 9c) and thus back-mutation of F312L to recover agonist potency was not pursued as it was deemed unlikely that the resultant receptor would be stable enough for NMR experiments.

α_1A_-AR-A4-active was labelled with ^13^C^ε^H_3_-methionine and 2D ^1^H-^13^C SOFAST-HMQC spectra acquired as above (Figure 5a,c). Overall the ligand-perturbed chemical shifts of the ^13^C^ε^H_3_-methionine resonances in α_1A_-AR-A4-active were similar to those in α_1A_-AR-A4 and α_1A_-AR-A4 (L312F), except for several key differences with the Met115^3.41^ and Met203^5.57^ resonances. We acquired spectra of four independent preparations of apo α_1A_-AR-A4-active (four biological replicates) and found that in the absence of bound ligand the data were not easily reproduced (Supplementary Figure 10a). The well resolved Met203^5.57^ varied between an intense peak, two peaks of similar intensity, or a peak of weak intensity. Met115^3.41^ persisted as a split peak, although the two components varied in intensity. Importantly, in the presence of the most potent inverse agonist, prazosin, the resonances of Met115^3.41^ and Met203^5.57^ were single peaks, and regardless of sample preparation, exhibited the same chemical shifts. The ^13^C^ε^H_3_ Met115^3.41^ resonance, as perturbed by prazosin, aligned approximately with the upfield component of the resonance for apo α_1A_-AR-A4-active (Supplementary Figure 10b). In contrast, upon titration with the neutral antagonist, silodosin, the peaks of Met115^3.41^ also collapsed to a single resonance with identical chemical shifts, but now aligned best with the downfield component of apo α_1A_-AR-A4-active (Supplementary Figure 10b). For the two partial inverse agonists, WB-4101 and phentolamine, the ^13^C^ε^H_3_ resonance of Met115^3.41^ was a single resonance, positioned midway between the ‘prazosin’ (upfield) and ‘silodosin’ (downfield) peaks (Figure 5d). These trends for ligand-efficacy were present in α_1A_-AR-A4 and α_1A_-AR-A4 (L312F), but were not as distinct as now observed for α_1A_-AR-A4-active, and notably the apo states for α_1A_-AR-A4 and α_1A_-AR-A4 (L312F) did not show two discrete peaks for Met115^3.41^. Such apo state sample-to-sample heterogeneity may suggest the presence of misfolded contaminants, but upon the addition of prazosin or silodosin each of these samples gave identical spectra (Supplementary Figure 10a), supporting the binding competency of the α_1A_-AR-A4-active samples. The diversity of apo state spectra likely reflects diversity of conformational states of similar free energy. The addition of agonist again resulted in, a single resonance for ^13^C^ε^H_3_ Met115^3.41^ that shifts upfield in ^1^H and downfield in ^13^C (Figure 5d). The trend in shifts of these resonances, however, suggests they follow in a linear manner evolving from the downfield (basal) signal of the apo state and reflects the selection of the active-like state.

**Figure 5.**
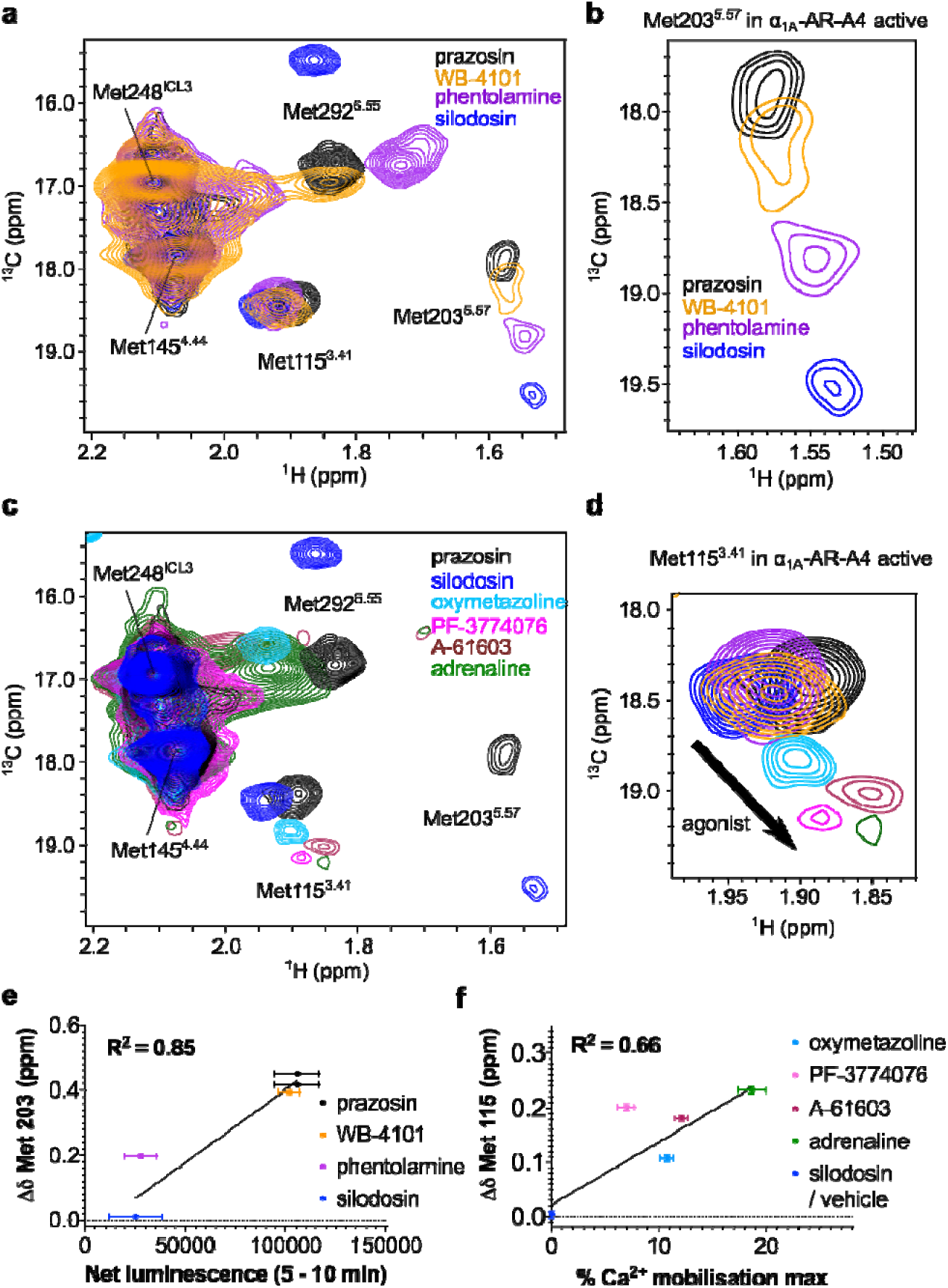
^1^H-^13^C SOFAST-HMQC spectra of α_1A_-AR-A4-active. (a) Overlay of 2D ^1^H-^13^C SOFAST-HMQC spectra of [^13^C^ε^H_3_-Met] α_1A_-AR-A4-active bound to prazosin (black, inverse agonist), WB-4101 (yellow, inverse agonist), phentolamine (purple, inverse agonist) and silodosin (blue, neutral antagonist). (b) Close-up of the Met203^5.57^ resonance. (c) Overlay of 2D ^1^H-^13^C SOFAST-HMQC spectra of [^13^C^ε^H_3_-Met] α_1A_-AR-A4-active bound to prazosin (black, inverse agonist), silodosin (blue, neutral antagonist), oxymetazoline (cyan, partial agonist), PF-3774076 (magenta, partial agonist), A-61603 (maroon, full agonist), and adrenaline (green, full agonist). (d) Close-up of the Met115^3.41^ resonance. (e) Linear regression analysis of the average chemical shift differences (Δδ) for the ^13^C^ε^H_3_ of Met203^5.57^ in α_1A_-AR-A4-active when bound to prazosin (black circles), WB-4101 (orange circles), and phentolamine (purple circles) compared to silodosin (blue circles) and the increase in luminescence seen in the NanoBit assay with α_1A_-AR-A4-active expressing COS-7 cells treated with the same antagonist (from Supplementary Figure 11a). Testing the resultant equation against the null hypothesis of a slope of zero resulted in a P value of 0.0041 (f) Linear regression analysis of the average chemical shift differences (Δδ) for the ^13^C^ε^H_3_ of Met115^3.41^ in α_1A_-AR-A4-active when bound to oxymetazoline (cyan circles), PF-3774076 (pink circles), A-61603 (dark red circles), adrenaline (green circles) and silodosin (blue circles) and the efficacy of each agonist in triggering Ca^2+^ mobilization in α_1A_-AR-A4-active expressing COS-7 cells (from Supplementary Figure 8b-e). Testing the resultant equation against the null hypothesis of a slope of zero resulted in a P value of 0.0154 In (e) and (f) Δδ are plotted for two independent titrations of prazosin, silodosin and oxymetazoline, and single experiments for other ligands. In (a-d) spectra were acquired on ∼50 µM α_1A_-AR-A4-active dissolved in 0.02-0.1% DDM micelle, pH 7.5 and 25 °C. Average chemical shift differences (Δδ) were normalised using the equation Δδ=[(Δδ_1H_)^2^+(Δδ_13C_/3.5)^2^]^0.5^ and errors were calculated by the formula [Δδ_1H_*R_1H_+Δδ_13C_*R_13C_/(3.5)^2^]/Δδ, where R_1H_ and R_13C_ are the digital resolutions in ppm in the ^1^H and ^13^C dimensions respectively (Kofuku et al., 2012).

A major difference between the spectra of α_1A_-AR-A4 and α_1A_-AR-A4-active, however, was significantly increased line broadening of the Met203^5.57^ signal of α_1A_-AR-A4-active in the apo state (Supplementary Figure 10a) and when bound to antagonists (Figure 5b). This broadening suggests that the DRY motif near the G protein-binding site of α_1A_-AR-A4-active is more dynamic compared to α_1A_-AR-A4, consistent with a receptor that more readily transitions between inactive and active-receptor states. Importantly, similar to α_1A_-AR-A4 and α_1A_-AR-A4 (L312F) variants, the ^13^C^ε^H_3_ of Met203^5.57^ shows an efficacy-dependent linear ^13^C^ε^H_3_ chemical shift change in the presence of inverse agonist and neutral antagonist, trending to an upfield ^13^C position (χ3 of ±gauche) for the more potent inverse agonist (Figure 5a,b). Unexpectedly, the addition of silodosin (neutral antagonist) resulted in a significant ^13^C downfield shift to near 19.5 ppm consistent with a trans χ3 angle for Met203^5.57^. Furthermore, similar to α_1A_-AR-A4, the Met203^5.57^ ^13^C^ε^H_3_ resonance of α_1A_-AR-A4-active was near completely broadened in the presence of agonists.

NanoBiT G protein activity assays were performed on COS-7 cells expressing α_1A_-AR-A4-active to determine relative inverse agonist efficacies. The inverse agonists reduced basal Gq activity in α_1A_-AR-A4-active in a similar way to WT α_1A_-AR expressing cells (Figure 4a and Supplementary Figure 11a). The net luminescence change induced by each inverse agonist at α_1A_-AR-A4-active expressing cells correlated well, in a linear fashion, with the ^13^C^ε^H_3_ chemical shift changes of Met203^5.57^ that each ligand induced at purified α_1A_-AR-A4-active, when analysed over two separate time periods (Figure 5e and Supplementary Figure 11b). Interestingly, the agonist-induced chemical shift changes of ^13^C^ε^H_3_ Met115^3.41^ showed a linear correlation with the efficacy of each agonist in Ca^2+^ mobilization assays on α_1A_-AR-A4-active expressing COS-7 cells (Figure 5f) although the partial agonist PF-3774076 was a notable outlier. Overall these cell-based assays with the ligand efficacy-correlated chemical shift changes of Met115^3.41^ and Met203^5.57^ clearly demonstrate that a conformational selection mechanism underlies receptor function in cells.

## Discussion

Recent spectroscopic studies have demonstrated that different classes of GPCR ligands distinctly alter the population of receptor states within the GPCR conformational equilibrium(Shimada et al., 2018). GPCR conformational changes are driven by defined structural changes in the microswitches (Ahuja and Smith, 2009; Deupi and Standfuss, 2011; Latorraca et al., 2017; Trzaskowski et al., 2012) (Figure 1) and, thus, how particular ligands affect the GPCR microswitch states likely underlies their pharmacological output as inverse, partial, full or biased agonists. Observing these effects, however, remains challenging. α_1A_-AR was one of the first GPCRs to be cloned and pharmacologically characterised (Cotecchia et al., 1988) and is clinically targeted with agonists as nasal decongestants and antagonists for hypertension and BPH. Despite the importance of this receptor there are currently no three-dimensional structures of α_1A_-AR, reflecting the inherent instability of this protein. Here, we demonstrate that prototypical ligands modulate the conformational equilibrium, as measured at the microswitches, of α_1A_-AR in defined and predictable ways by ^13^C^ε^H_3_-methionine labelling α_1A_-AR variants and monitoring the ^1^H and ^13^C chemical shifts of these methyl resonances in the presence of ligands of different efficacy,.

It is well accepted that the NMR signals of methionine methyl groups are sensitive to the local environment and the conformation of the methionine side chain (Kofuku et al., 2012; Nygaard et al., 2013). Many of the conclusions made in this study rely on Met115^3.41^, a probe for the conformation of the transmission switch, and Met203^5.57^ as a probe of the DRY motif that signifies intracellular TMD rearrangements for G-protein binding. In our model of α_1A_-AR, Met203^5.57^ sits over Tyr125^3.51^ of the DRY motif but is distant from Arg124^3.50^ which is expected to undergo significant rotameric changes within this motif (Carpenter and Tate, 2017) (Figure 1d). Met115^3.41^ is sequential to the Ile114^3.40^ in TM3 but it points away and is distant to transmission switch residue Phe281^6.44^ located on TM6 that is expected to undergo significant rotameric changes (Figure 1c). While in the thermostabilized inactive α_1A_-AR-A4 mutant the residues of the transmission switch and DRY motif are retained, the asparagine of a third microswitch, the NPxxY motif, is mutated to lysine, which likely forms a salt bridge with Asp72^2.50^ to lock this switch in an inactive state. The transmission switch and NPxxY motif are proximal to each other and therefore Met115^3.41^, while distant to NPxxY, is likely to be sensitive to conformational changes involving both switches. In the reported active-state GPCR structures, three conserved residues (Arg^3.50^ of the DRY motif, Tyr^7.53^ of the NPxxY motif and Tyr^5.58^) adopt near identical positions and connect these microswitches through water-mediated hydrogen bonds(Carpenter and Tate, 2017; Manglik and Kruse, 2017). Furthermore, in our model of active α_1A_-AR which is based on structures of β_2_-AR, Arg124^3.50^ of the DRY motif is in contact with Tyr326^7.53^ of the NPxxY motif (Figure 1d).

In our NMR experiments for all ligands the ^13^C^ε^H_3_ group of both Met115^3.41^ and Met203^5.57^ show significant directional chemical shift and line-width changes that are correlated with ligand efficacy, not affinity. As a distinct peak is observed for the addition of each ligand the chemical shift likely reflects an average population exchanging on a fast to intermediate timescale. The chemical shift differences, however, reflect a shift in the equilibrium, and specifically for the ^13^C^ε^H_3_ of Met203^5.57^, from a χ3 of a gauche-trans average (inverse agonist) towards a trans (agonist) average (Figure 6a). An NMR study using ^15^N-labelled, thermostabilised β_1_AR observed substantial ligand efficacy-correlated backbone chemical shift changes for V226^5.57^, which is in the same position as Met203^5.57^ in α_1A_-AR (Isogai et al., 2016). The authors speculated that these changes were caused by TM5 bending towards the active receptor state (Isogai et al., 2016), an idea that may also apply to α_1A_-AR and other GPCRs. Here, the linear chemical shift changes of Met203^5.57^, and to a lesser degree Met115^3.41^, in response to ligands of different efficacy is strong evidence that agonists activate α_1A_-AR via a conformational selection mechanism. The line broadening of Met115^3.41^ and Met203^5.57^ upon agonist binding supports an efficacy-driven shift in dynamics, and thereby the equilibrium of conformational states, communicated allosterically by the microswitches and sensed by these methionine residues (Figure 6).

**Figure 6.**
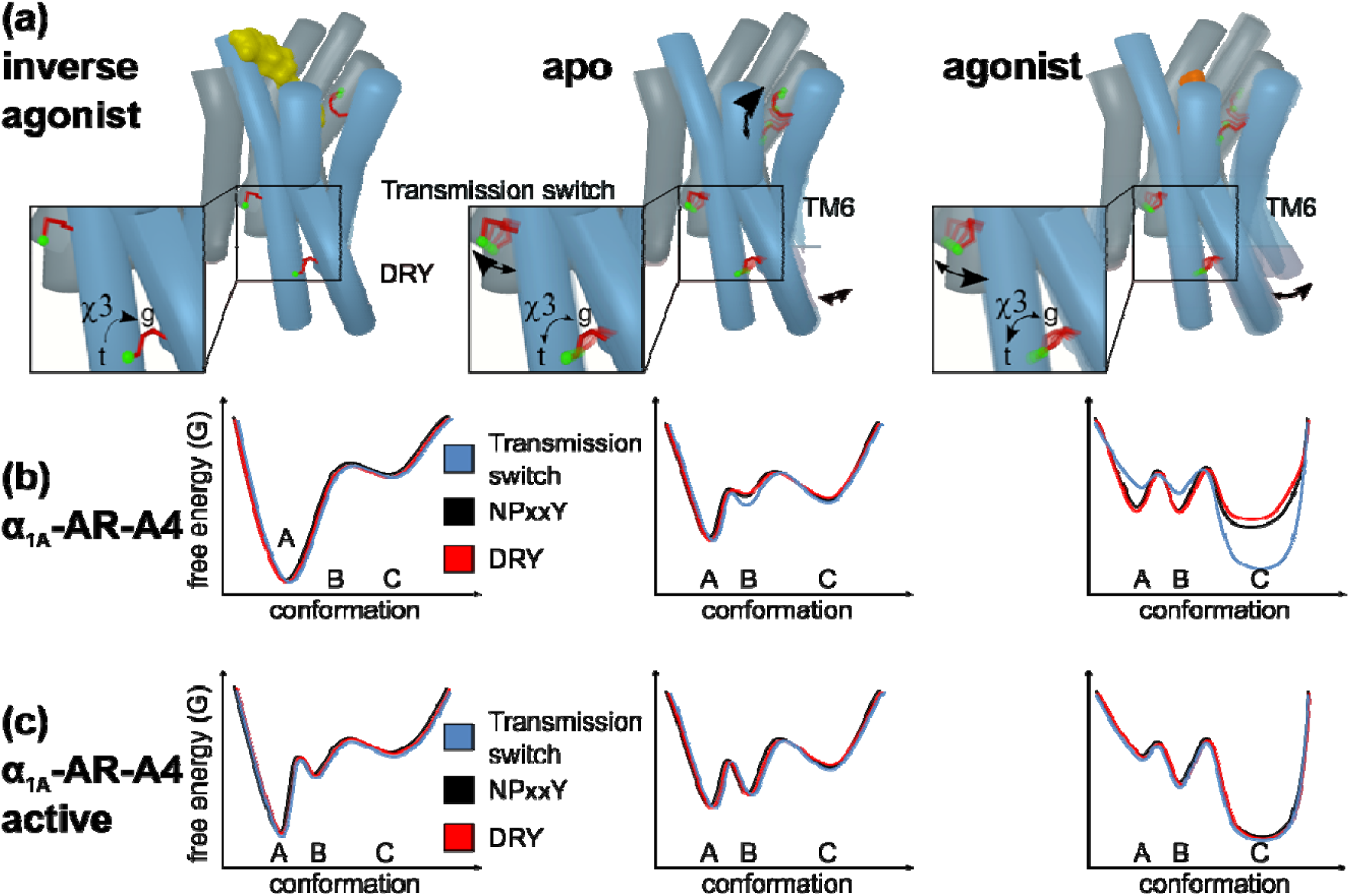
How ligands modulate the conformational landscape of the α_1A_-AR microswitches. (a) Cartoon representations of α_1A_-AR in the inverse agonist-bound, apo, and agonist-bound states. The three probe methionines, Met292^6.55^ (binding site), Met115^3.41^ (transmission switch) and Met203^5.57^ (DRY microswitch) are highlighted with red sticks and the labeled methyl group in green. The arrows labeled χ3 illustrate the ligand induced changes to the equilibrium between the trans (t) and gauche (g) χ3 dihedral angle of Met203^5.57^. Other arrows indicate how different ligands alter the conformation equilibria of Met292^6.55^, Met115^3.41^ and TM6. Hypothetical free energy landscape diagrams of the three microswitches in (b) the inactive α_1A_-AR-A4 receptor compared to (c) α_1A_-AR-A4-active. (A) indicates the proposed inactive states, (B) represents basal states, and (C) represents active states of the microswitches.

For our signalling incompetent receptors, α_1A_-AR-A4 and α_1A_-AR-A4 (L312F), the NPxxY microswitch has been mutated (N322K) in a way that would bias the NPxxY switch towards inactive states (K322 - D72 salt bridge). A consequence of this mutation is that for α_1A_-AR-A4 (L312F) the DRY motif probe, Met203^5.57^, gives relatively intense chemical shifts in the apo and antagonist-t-bound states (conformational equilibrium biased towards inactive states) (Figure 6b). On restoration of the NPxxY microswitch in α_1A_-AR-A4-active however, the Met203^5.57^ chemical shifts broaden and shift towards the agonist bound position (as defined for α_1A_-AR-A4 (L312F)), even with neutral antagonist bound. Therefore, restoring the NPxxY microswitch enables the DRY motif of α_1A_-AR-A4-active to more readily sample active-like states. The full trans (19.5 ppm in ^13^C) populated by silodosin indicates that we may observe the full-active state, although previous studies showed that the full-active state is only populated in the presence of both agonist and nanobody (Manglik et al., 2015; Nygaard et al., 2013; Solt et al., 2017; Sounier et al., 2015; Xu et al., 2019). Our two methionine probes, Met115^3.41^ and Met203^5.57^, retain similar ligand-induced behaviour in α_1A_-AR-A4-active compared to α_1A_-AR-A4 and α_1A_-AR-A4 (L312F), where the latter are both essentially inactive. The chemical shift and line-broadening trends of Met115^3.41^ and Met203^5.57^ suggest that the transmission switch and DRY motif, in the presence of an inactive NPxxY motif, can independently adopt conformations representative of active and inactive states. To fully adopt the conformational signatures of an active receptor, a functional NPxxY motif is required (in α_1A_-AR-A4-active) thus increasing the dynamics of the transmission switch and DRY motif, suggesting that the interdependence of the three microswitches is a consequence of their dynamic nature, and that this is required for full receptor function.

While the striking linear chemical shift dependence for the ^13^C^ε^H_3_ of Met203^5.57^ on ligand efficacy is consistent with a smooth change in equilibria from inactive to active, linear trends for ^13^C^ε^H_3_ of Met115^3.41^ were less clear. In the inactive variants α_1A_-AR-A4 and α_1A_-AR-A4 (L312F) the resonance for apo, inverse agonist and neutral antagonist shows little variation, but clear chemical shift changes and broadening are observed for agonists. The most distinct changes for the ^13^C^ε^H_3_ of Met115^3.41^ were for α_1A_-AR-A4-active, where in the apo-state two peaks were consistently observed, suggesting slow exchange (> millisecond) between two distinct states, to which we attribute to restoring the NPxxY microswitch. On the basis of chemical shift, we propose that the upfield peak of apo α_1A_-AR-A4-active represents fully inactive receptor (state A in Figure 6), expected for full inverse agonists, and the downfield peak with a basal state receptor that is stabilized by the neutral antagonist (state B in Figure 6). This downfield ‘basal’ peak shows approximate linear efficacy-dependent chemical shift changes with agonist titrations. Therefore, the ^13^C^ε^H_3_ of Met115^3.41^ reflects three states, (inverse agonist) inactive, an intermediate (basal) and active states (state(s) C in Figure 6), where the latter progressively shift from partial to full agonist states. Interestingly, in a ^13^C^ε^H_3_-methionine labelled study on the M2R, Met112^3.41^, which is equivalent to Met115^3.41^ in α_1A_-AR, did not display efficacy-dependent chemical shift changes. In the presence of ligands, however, M2R Met112^3.41^ was resolved as two separate resonances, consistent with a slow exchanging microswitch(Xu et al., 2019) and may highlight some differences between how different rhodopsin family GPCRs function.

In this NMR study, by starting with a signalling incompetent variant of α_1A_-AR and subsequently restoring signalling activity through back mutations, we were able to study the functional dynamics of the key GPCR microswitches and how different ligands modulate this. Our NMR data for the transmission and DRY microswitches revealed ligand efficacy-dependent changes to the microswitch conformational equilibria, supporting a conformational selection mechanism for α_1A_-AR modulation. This and the agonist-driven line broadening for both microswitches suggest similar mechanistic actions on the different microswitches, supporting ligand-driven allosteric communication between the microswitches. MD simulations (Dror et al., 2011) of β_2_-AR suggest that these microswitches behaved independently of each other, with only loose allosteric coupling. While this may be true over the relatively short timescales of MD, we believe that over the course of an NMR experiment such loose allosteric coupling culminates in significant coupled shifts to the microswitch conformations and that this is likely how ligands modulate GPCR signalling in cells.

### Data Availability

All data that support the conclusions are included in the published paper and its supplementary information, or are available from the authors on request.

## Acknowledgements

We thank Dr Fabian Bumbak (The Florey Institute of Neuroscience and Mental Health) for assistance with optimising the expression and purification of receptor samples; Prof. Dmitry Veprintsev (University of Nottingham) and Dr Franziska Heydenreich (Stanford University) for suggesting back mutations for the generation of α_1A_-AR-A4-active; Prof. Asuka Inoue (Tohoku University) for supplying plasmids for the G protein activity assays; Sharon Layfield (The Florey Institute of Neuroscience and Mental Health) and Dr Ashish Sethi (The University of Melbourne) for assistance with cell-based assays and NMR data analysis respectively; Dr David Chalmers (Monash University) and the Monash University Medicinal Chemistry Computational Chemistry Facility for the assistance with computational modelling; The Bio21 NMR facility for access to spectrometers. This work was supported by NHMRC project grants 1081801 (D.J.S), 1081844 (P.R.G, D.J.S, R.A.D.B) and 1141034 (D.J.S, P.R.G, R.A.D.B). D.J.S. is an NHMRC Boosting Dementia Research Leadership Fellow.

## Author contributions

FJW performed cloning, mutagenesis, protein expression and purification, thermostability assays, acquisition and analysis of NMR data; LMW competition binding assays and saturation binding assays; AAR intracellular Ca^2+^ mobilization assays; AG, MK, and RADB NanoBiT G protein activity assays; TV computational modelling; ARW IP_1_ assays. DJS, MDWG and PRG conceived the experiments and with FJW analysed, prepared figures and wrote the manuscript. All authors contributed to the editing of the manuscript.

## Methods

### α_1A_-AR constructs

The α_1A_-AR-A4 variant is a thermostabilised human α_1A_-AR, containing 15 stabilising mutations (Yong et al., 2018). Met80^2.58^ in the spectra was introduced through the stabilisation process in lieu of the naturally occurring amino acid leucine. As compared to wild type, the carboxyl termini of the α_1A_-AR-A4 variant was modified by truncation at Ser351 and addition of a deca-His tag to facilitate purification (Supplementary Figure 1). For expression, the α_1A_-AR variants sequences were sub-cloned into the pQE30 derived vector, pDS15, with a maltose-binding protein (MBP) and a methionine-free monomeric ultra-stable green fluorescent protein (-Met-muGFP) (Scott et al., 2018) attached respectively to the N- and C-termini of the receptor via HRV 3C protease cleavage sites. For the purpose of Kingfisher binding assays, α_1A_-AR variants were sub-cloned into a similar vector, pDS11, in which muGFP was replaced with mCherry since the excitation and emission wavelengths of fluorescent QAPB were overlapping with those of GFP. The final sequence of α_1A_-AR-A4 after purification (with residues left from HRV 3C cleavage) is: GPGSVFLSGNASDSSNSIQPPAPVNISKAILLGVILGGIILFGVLGNILVILSVACHRHLHSVTHYYVVYLAVADLLLTSTVMPFSAIYEVLGYWAFGRVFCNIWAAVDVLCCTASIMGLCIISIDRYIAVSYPLRYPTIVTQRRALMALLCVFALSLVISIGPLFGWRQPAPVDETICQINEEPGYVLFSALGSFYLPLAIILVMYCRVYVVAKRESRGLKSGLKTDKSDSEQVTLRIHRKNAPAGGSGMASAKTKTHFSVRLLKFSREKKAAKTLGIVVGCFVLCWLPFFLVMPIGSFFPDFKPSETVFKIVLWLGYLNSCIKPIIYLCYSQEFKKAFQNVLRIQCLCRKQSASHHHHHHHHHHGTRSLRGGLEVLFQ

In the α_1A_-AR-A4 (L312F) variant one stabilising mutation L312 was reversed to phenylalanine to improve the affinity of ligands compared to α_1A_-AR-A4. α_1A_-AR-A4-active (Y67N, K322N) is a signalling competent variant in which the two stabilizing mutations Y67 and K322 were reverted to the wild-type asparagines. α_1A_-AR-A4 was used for NMR assignment, where each methionine was substituted to either leucine or isoleucine. All mutations were introduced through site-directed mutagenesis using PrimeSTAR DNA polymerase (TaKaRa).

### α_1A_-AR expression

All α_1A_-AR variants were expressed in *E. coli* C43 (DE3) cells (Lucigen, Middleton, WI). For ^13^C^ε^H_3_-methionine labelled expressions, 5 mL LB pre-culture containing 100 mg/L ampicillin and 1% (w/v) glucose was inoculated with a single colony of C43 cells freshly transformed with the expression plasmid and incubated at 37 °C, 225 rpm for approx. 8 h. 2 mL of the LB day culture was centrifuged (1700 rcf, 22 °C, 5 min) and the pellet was used to inoculate 100 mL of a defined minimal medium (M1 medium) (Bumbak et al., 2018) as overnight pre-culture. 10 mL of the overnight pre-culture was used to inoculate 500 mL of M1 medium in 2 L flasks. The expression cultures were incubated at 37 °C, 225 rpm to reach OD_600_ of 0.6, at which point 50 mg/L ^13^C^ε^H_3_-methionine (Cambridge Stable Isotopes) was added along with 100 mg/L of each lysine, threonine, phenylalanine, and 50 mg/L of each leucine, isoleucine and valine. The flasks were transferred to 20 °C and left shaking for 15 min prior to inducing protein expression with 250 µM isopropyl β-D-1-thiogalactopyranoside (IPTG). After overnight expression (15-18 h, 20 °C, 225 rpm), the culture was harvested by centrifugation (2600 rcf, 4 °C, 15 min). The final pellet was snap frozen in liquid nitrogen and stored at -80°C. For unlabelled expression 5 mL of LB pre-culture was used to inoculate 500 mL 2YT medium containing 100 mg/L ampicillin and 0.4% (w/v) glucose. At OD_600_ of 0.6 the culture was chilled on ice for 2 min prior to inducing protein expression with 250 µM IPTG overnight expression and harvesting were carried out as described above.

### α_1A_-AR purification

The frozen cell pellet was thawed at room temperature for 30 min. 10 mL pellet was gently resuspended in 40 mL ice-cold solubilisation buffer [25 mM HEPES, pH 7.5, 200 mM NaCl, 10% glycerol, 1% DDM (Anatrace), 0.12% CHS (cholesterol hemi succinate, Anatrace), 0.6% CHAPS (Sigma), 50 mg lysozyme, 5 mg Dnase, one tablet of EDTA free complete protease inhibitor cocktail (Roche), 0.2-0.4 mM PMSF (phenylmethylsulfonyl fluoride)] and incubated on a turning wheel for 30 min at 4 °C. The cell membranes were then disrupted by sonication device (Diagenode Bioruptor Plus, high power, 10s on/20s off for 30 cycles) followed by another 1 h incubation at 4 °C on the turning wheel. The cell debris was removed by centrifugation (12,000 rcf, 4 °C, 40 min) and the supernatant was filtered using a 45 µm Durapore syringe filter (Merck Millipore). The cleared cell lysate was incubated with 3 mL Talon metal affinity resin pre-equilibrated with 45 mL equilibrium buffer (20 mM HEPES, pH 7.5, 300 mM NaCl, 10% glycerol, 0.05% DDM). After 1.5 h incubation at 4 °C, the resin retaining the receptor was washed three times with washing buffer 1 (20 mM HEPES, pH 7.5, 500 mM NaCl, 10% glycerol, 0.05% DDM) and then the full-length protein was eluted by 30 mL elution buffer (20 mM HEPES, pH 7.5, 300 mM NaCl, 10% glycerol, 0.05% DDM, 250 mM Imidazole). The eluate was concentrated down to 0.5-1 mL using a 100 kDa cut-off centrifugal filter device (Amicon Ultra, Millipore). Imidazole was removed by using a PD10 desalting column (GE Healthcare). Cleavage of fusion proteins from the receptor was carried out overnight at 4 °C by adding 100 mM Na_2_SO_4_, 1 mM TCEP and 300 pmol GST-tagged HRV 3C protease (made in house).

The cleaved mixture was incubated for 1 h with 2 mL of pre-equilibrated Talon resin. The resin was washed using 30 mL washing buffer 2 (20 mM HEPES, pH 7.5, 300 mM NaCl, 10% glycerol, 0.05% DDM, 30 mM Imidazole) and the receptor was eluted by 20 mL elution buffer. The eluate was concentrated down to 450 µL by 30 kDa cut-off centrifugal filter device (Amicon Ultra, Millipore) and it was loaded onto a Superdex 200 10/300 increase column (GE healthcare) equilibrated with SEC buffer (50 mM sodium phosphate, pH 7.5, 100 mM NaCl, 0.02% DDM). Size exclusion chromatography (SEC) was carried out at a flow rate of 0.5 mL/min. The peak fractions containing receptor were pooled and concentrated down to 100 µL using a 30 kDa cut-off centrifugal filter device (Amicon Ultra, Millipore). The sample buffer was exchanged twice to NMR buffer (50 mM sodium phosphate, pH 7.5, 100 mM NaCl, 99.9% D_2_O). Yields were generally between 0.5-1 mg receptor per litre of expression culture. Protein concentration was measured by BCA protein assay (Pierce, ThermoFisher).

### NMR spectroscopy

NMR samples were prepared to 130 µL at 40-60 µM receptor in a 3 mm Shigemi NMR tubes (Shigemi Inc, Allison Park, PA). Ligands were added at saturating concentrations that were 2 mM adrenaline for α_1A_-AR-A4 and α_1A_-AR-A4 (active), 400 µM prazosin, and 1 mM of other ligands to all mutants (supplementary Table 1). Adrenaline was supplemented with 1 mM of the anti-oxidant ascorbic acid. Samples containing low affinity agonists (phenylephrine and adrenaline) were recycled via competition with high affinity ligands, exchange was judged via the chemical shift of the Met292^6.55^ resonance. Experiments on α_1A_-AR-A4 and α_1A_-AR-A4-L312F, apo and all ligands were performed at least twice on independent receptor samples (biological replicates), except WB-4101 and phentolamine which were acquired once. Experiments on apo α_1A_-AR-A4-active and bound to prazosin, silodosin, and oxymetazoline were performed at least twice on independent receptor samples, and for other ligands were performed once.

All NMR spectra were collected at 25 °C on an 800-MHz Bruker Avance II spectrometer equipped with a triple resonance cryoprobe. 2D ^1^H-^13^C SOFAST-HMQC (Schanda et al., 2005) spectra were recorded by excitation with a 2.25 ms PC9 120 degree ^1^H pulse and refocusing with a 1 ms r-SNOB shaped 180 degree ^1^H pulse. The spectral widths were set to 12 ppm and 25 ppm for ^1^H and ^13^C dimensions respectively. For the spectra recorded for α_1A_-AR-A4 variant (Figure 2 and Supplementary Figure 3), 1024 x 128 complex points were recorded with a 25% Poisson-gap sampling schedule and 2048 scans; an acquisition time of 8.5 h. For the other spectra, 1024 x 200 complex points were recorded with either traditional or 60% Poisson-gap sampling schedule and 368 scans resulting in acquisition times of 10 h and 6 h respectively. Spectra were reconstructed with compressed sensing using qMDD and processed using NMRpipe (Delaglio et al., 1995) where data were multiplied by cosine bell functions and zero-filled once in each dimension. Spectra were analysed in NMRFAM-Sparky(Lee et al., 2015) (Goddard, T.D. and Kneller, D.G, University of California, San Francisco).

The average chemical shift differences, Δδ, were normalised using the equation Δδ=[(Δδ_1H_)^2^+(Δδ_13C_/3.5)^2^]^0.5^. The error values were calculated by the formula [Δδ_1H_*R_1H_+Δδ_13C_*R_13C_/(3.5)^2^]/Δδ, where R_1H_ and R_13C_ are the digital resolutions in ppm in the ^1^H and ^13^C dimensions respectively (Kofuku et al., 2012).

### Saturation and Competition binding assays

1 nmol purified full-length α_1A_-AR variant (mCherry attached) was resuspended in 10 mL assay buffer (20 mM HEPES, pH 7.5, 100 mM NaCl, 0.02% DDM) and immobilized onto 200 µL of Dynabeads (Streptavidin T1) for 30 min at 4 °C. 100 µL of the suspension containing beads with immobilized receptor was aliquoted to a 96-DeepWell plate from which the beads transferred to another 96-DeepWell plate containing 100 µL ligand solution using a KingFisher Flex magnetic particle processor. For saturation binding, immobilized receptors in each well were incubated with 100 µL assay buffer containing increased concentration (0, 3.125, 6.25, 12.5, 25, 50, 100, 200 nM) of QAPB (Quinazoline Piperazine Bodipy) for 2 h at 22 °C. Nonspecific binding was determined by repeating the experiment in the presence of 10 µM of prazosin. For competition binding, immobilized receptors were incubated with 100 µL of assay buffer containing 10 nM QAPB with the addition of ligands at various concentrations, as shown in the Supplementary Figures 5 and 8, for 2 h at 22 °C. Immobilised receptors were subsequently washed with 200 µL of assay buffer and resuspended in 100 µL assay buffer. 90 µL of the final beads solution was transferred to a 96-well Greiner Bio-One nonbinding black plate. Fluorescence of bound QAPB was measured using a POLARstar OMEGA plate reader (BMG Labtech, Ortenburg, Germany) and normalised to mCherry fluorescence which was detected simultaneously. Data represent the mean ± standard deviation (SD) of three independent biological replicate experiments each performed in duplicate technical measurements. To compare ligand binding affinities at α_1A_-AR-A4 (L312F) and α_1A_-AR-A4-active of to α_1A_-AR-A4, raw data from our previously published paper (Yong et al., 2018), were reanalysed and presented in Supplementary Table 1.

### Thermostability assay

1 nM purified full-length α_1A_-AR-A4 or α_1A_-AR-A4-active (mCherry attached) was prepared in base buffer (20 mM HEPES, 100 mM NaCl, 0.1% DDM). To measure thermostability of receptors in the apo-state, 100 µL of receptor solution was aliquoted into 24 wells of a 96-well PCR plate. 10 of the 12 duplicates were heated in gradient temperatures for 30 min and the two remaining duplicates were left at 4 °C for normalisation. After thermo-treatment, the receptors were transferred to a KingFisher 96-DeepWell plate containing 2 µL paramagnetic Dynabeads per well (streptavidin T1, ThermoFisher Scientific). The following few steps were automatically performed by using a KingFisher 96 magnetic particle processor. The receptor was firstly incubated with magnetic beads for 30 min at 4 °C. Then, magnetic beads were transferred to another 96-DeepWell plate containing 100 µL ligand solution (20 mM HEPES, 100 mM NaCl, 0.1% DDM, 100 nM QAPB). The non-specific binding was determined by competing QAPB with 100 µL prazosin. After 1.5 h incubation, immobilised receptors were subsequently washed with 200 µL of assay buffer and resuspended in 100 µL assay buffer. 90 µL of the final beads solution was transferred to a 96-well Greiner Bio-One nonbinding black plate. Fluorescence of bound QAPB was measured using a POLARstar OMEGA plate reader (BMG Labtech, Ortenburg, Germany) and normalised to mCherry fluorescence which was detected simultaneously. To measure the thermostability of α_1A_-AR variants in the presence of ligand, receptors were preincubated with 100 nM QAPB for 1 h on ice prior to be heated at varying temperatures. The remaining steps were carried out as described for apo state thermostability assay. Data represent the mean ± SD of three independent biological replicate experiments each performed in duplicate technical replicate measurements.

### IP_1_ assay

Gαq/11 signalling assays were carried out using the IP-One HTRF® Assay Kit (Cisbio Bioassays, France) measuring inositol phosphate (IP_1_) using the manufacturer’s protocol. COS-7 cells were seeded at 25,000 cells per well in a 96-well plate and incubated overnight at 37 °C and 5% CO_2_ in Dulbecco’s modified Eagle medium (DMEM) (Gibco, Gaithersburg, USA) supplemented with 10% FBS (Scientifix Life, Melbourne, Australia), 1% L-Glutamine (Gibco) and 1% penicillin/streptomycin (Gibco). Cells were transfected with pcDNA3.1 constructs of WT or mutant α_1A_-ARs using Lipofectamine 2000 (Invitrogen, Carlsbad, USA) at 0.25 µg DNA per well. 24 h later, cells were stimulated with ligands for 2 h at 37 °C in 40 µL of Stimulation Buffer, then frozen at -80 °C. 14 µL of thawed sample were transferred to a white HTRF® 384-well Optiplate (PerkinElmer, Waltham, USA), incubated with development reagents in the dark for 1 h with shaking, and analysed by time-resolved fluorescence using a POLARstar OMEGA plate reader (BMG Labtech, Ortenburg, Germany). Data were analysed against the kit’s standard curve. Data represent mean ± SD of three independent biological replicate experiments each performed in triplicate technical replicate measurements, unless otherwise stated in the figure legends.

### NanoBiT G Protein Activity Assay

COS-7 cells grown in 10% fetal bovine serum (FBS), 1% L-Glutamine, 1% penicillin/streptomycin DMEM media were seeded at 250,000 cells per well on a six-well plate. Cells were then transiently co-transfected in the six-well plate, with 0.1 μg Gα_q_-LgBiT (GNAQ-11S) DNA, 0.5 μg Gβ-untagged (GNB1) DNA, 0.5 μg Gγ-SmBiT (114-GnG_2_) DNA, 0.2 μg Guanine Release Factor (RIC8A) DNA and 0.5 μg α_1A_-AR (or respective AR mutants) DNA using Lipofectamine 2000 transfection reagent as per the manufacturer’s instructions. The next day the cells were resuspended in Phenol-Red-free (PRF) DMEM media containing 10% FBS, 1% L-Glutamine, 1% penicillin/streptomycin, 25 mM HEPES and seeded at 50,000 cells per well to a white 96-well plate and incubated overnight. On the day of the assay, plates were pre-incubated with 10 μM Furimazine for 1 hour. Following incubation, raw luminescence counts in each well were measured every 12 sec over the course of the assay using a POLARstar Omega plate reader (BMG Labtech). Cells were treated with either vehicle or a saturating concentration of each ligand (50 nM for A-61603 and 100 nM for antagonists). Luminescence counts were plotted against time, with the final pre-incubation reading assigned as the zero-time point (time of vehicle/ligand addition). A baseline correction was then performed by subtracting the luminescence counts in the vehicle-treated samples from the ligand-treated samples which resulted in a time-course plot of ligand-induced luminescence counts. Initial raw luminescence counts were used as a readout of G protein expression levels. Data represent the mean ± standard error (SEM) of three independent biological replicate experiments each performed in triplicate technical replicate measurements, unless otherwise stated in the figure legends.

### Intracellular Ca^2+^ Mobilization Assays

COS-7 cells were seeded in 10 cm culture dishes at 3×10^6^ cells per dish and allowed to grow overnight at 37 °C, 5% CO_2_ in Dulbecco’s modified Eagle medium (DMEM) supplemented with 10% FBS, 1% L-Glutamine and 1% penicillin/streptomycin (Life Technologies, California, USA). The next day the cells were transfected with 30 µg of receptor DNA construct (pcDNA3.1 expression vector containing WT or mutant α_1A_-ARs) using 60 µl of Lipofectamine 2000 (Invitrogen) transfecting reagent per dish. The following day, cells were transferred to 96-well culture plates (5×10^4^ cells per well) and allowed to grow overnight. On the day of the experiment cells were washed twice with Ca^2+^ assay buffer [150 mM NaCl, 2.6 mM KCl, 1.2 mM MgCl_2_, 10 mM D-glucose, 10 mM HEPES, 2.2 mM CaCl_2_, 0.5% (w/v) BSA, and 4 mM probenecid, pH 7.4] and incubated in Ca^2+^ assay buffer containing 1 mM Fluo-4-AM for 1 h in the dark at 37 °C and 5% CO_2_. After two washes with Ca^2+^ assay buffer, fluorescence was measured for 1.5 min upon the addition of ligands in a Flexstation 3 (Molecular Devices, Sunnyvale, CA) using an excitation wavelength of 485 nm and emission wavelength of 520 nm. Data were normalized to the peak response elicited by 3 µM Ionomycin (Life Technologies). Data represent the mean ± SD of three independent biological replicate experiments each performed in triplicate technical replicate measurements, unless otherwise stated in the figure legends.

### Homology Modelling

Homology models of inactive- and active-state α_1A_-AR were built with I-TASSER (Zhang et al., 2015) using crystal structures of β_2_-AR as the templates. For inactive state models, the structure of β_2_-AR bound to the antagonist carazolol and the inactive-state stabilizing nanobody, Nb60 (PDB ID: 5JQH) (Staus et al., 2016) was used as a template. For the active state models, the crystal structure of a β_2_-AR-Gs protein complex bound to the agonist BI-167107 (PDB ID: 3SN6) (Rasmussen et al., 2011) was used as a template. The N- and C-terminal regions as well as the ICL3 regions, which have no sequence similarity to the template, were deleted from the model. Energy minimisation was performed using Minimize tool in Maestro version 11.7.012 (Schrödinger, Inc.) under OPLS 2005 (Siu et al., 2012) forcefield.

## Supplementary Figures and Tables

**Supplementary Figure 1.**
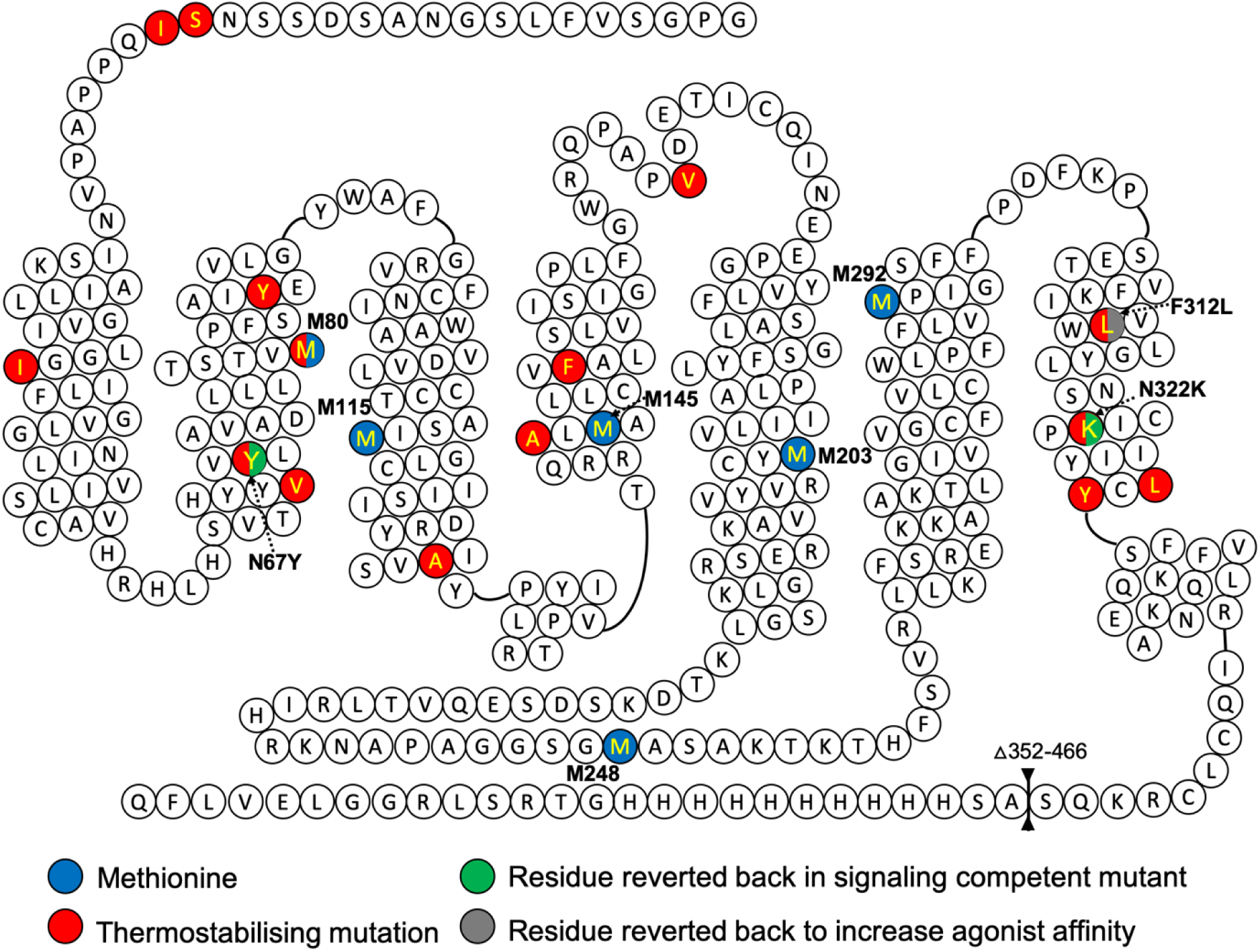
Secondary structure diagram of human α_1A_-AR-A4. Methionine residues and thermostabilisation mutations are labelled in blue and red, respectively. Two thermostabilisation mutations (N67Y, N322K) labelled in both red and green were reverted back to natural residues in making signaling competent construct α_1A_-AR-A4-active (Y67N, K322N). Residue labelled in grey is critical for ligand binding, α_1A_-AR-A4 (L312F) construct was made to rescue affinities of agonists tested in this study.

**Supplementary Figure 2.**
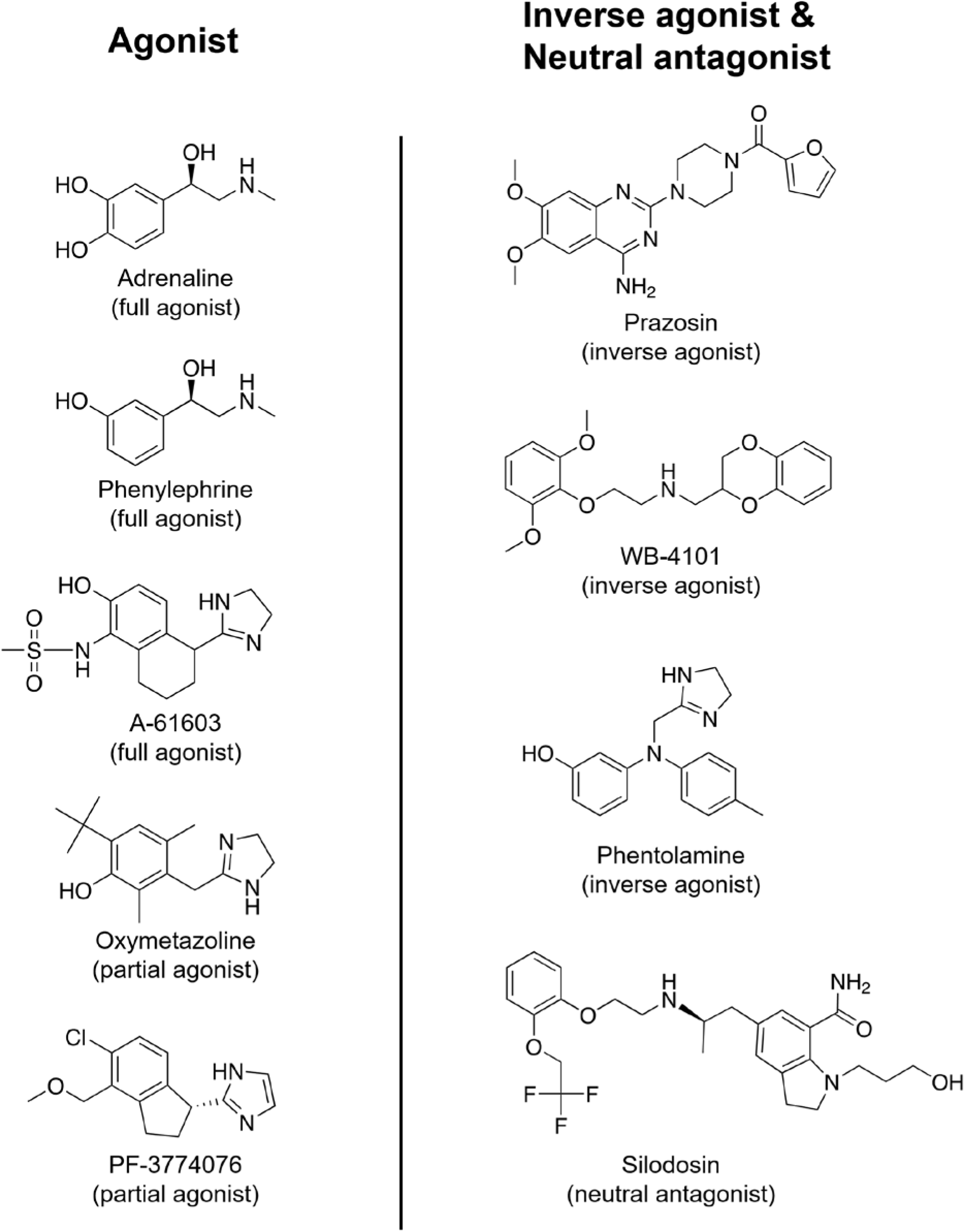
Chemical structures of ligands used in this study.

**Supplementary Figure 3.**
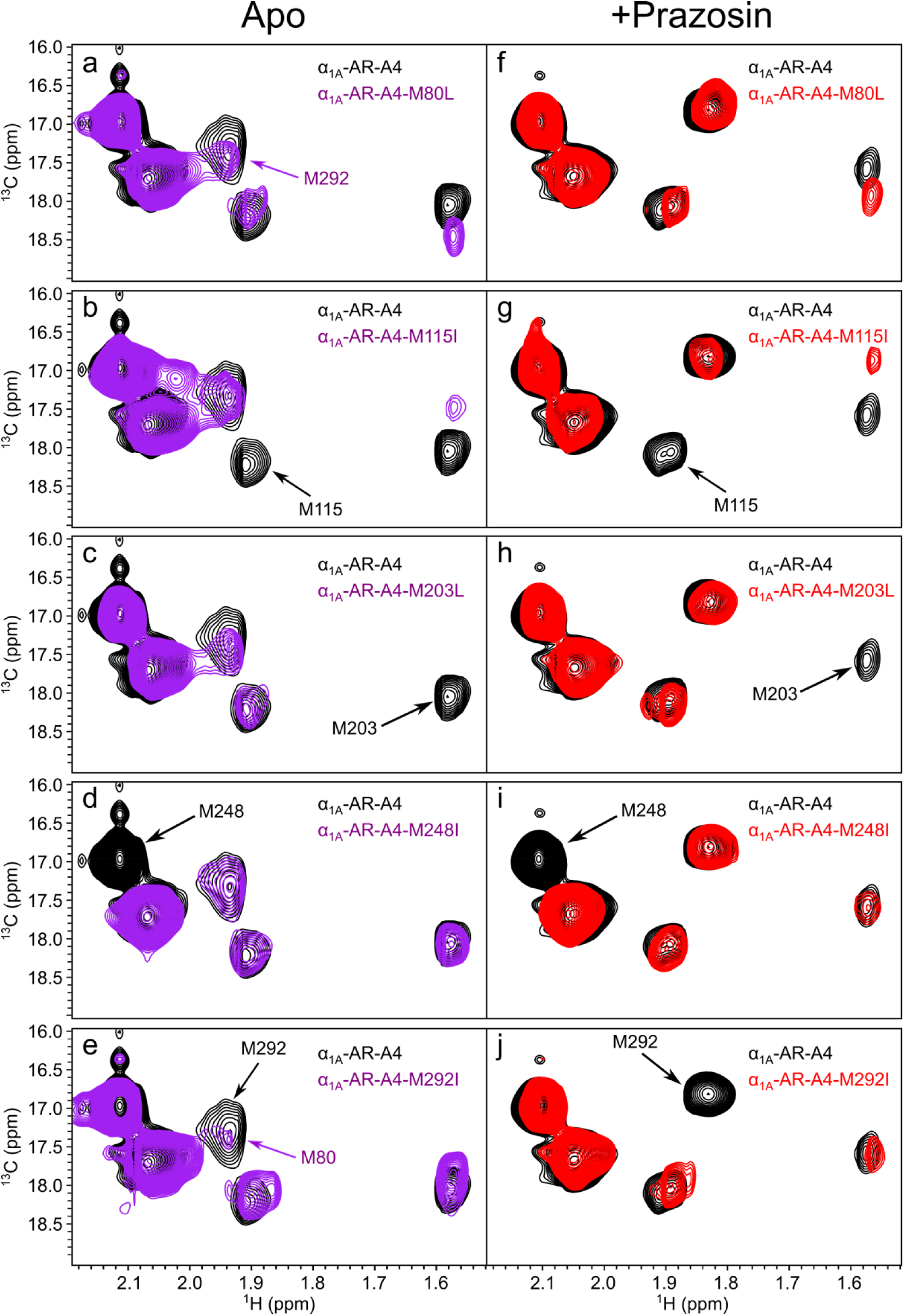
Assignment of ^13^C methyl labelled methionine residues in α_1A_-AR-A4. Five methionine residues in α_1A_-AR-A4 were individually mutated to either Leucine or Isoleucine, M80L (a,f); M115I (b,g); M203L (c,h); M248I (d,i); M292I (e,j). The ^1^H-^13^C SOFAST HMQC spectra of five α_1A_-AR-A4 mutants were collected in apo state (a-e, purple) and prazosin-bound state (f-j, red). Spectra of all mutants overlay with the spectrum of α_1A_-AR-A4 in the apo or prazosin-bound state (black).

**Supplementary Figure 4.**
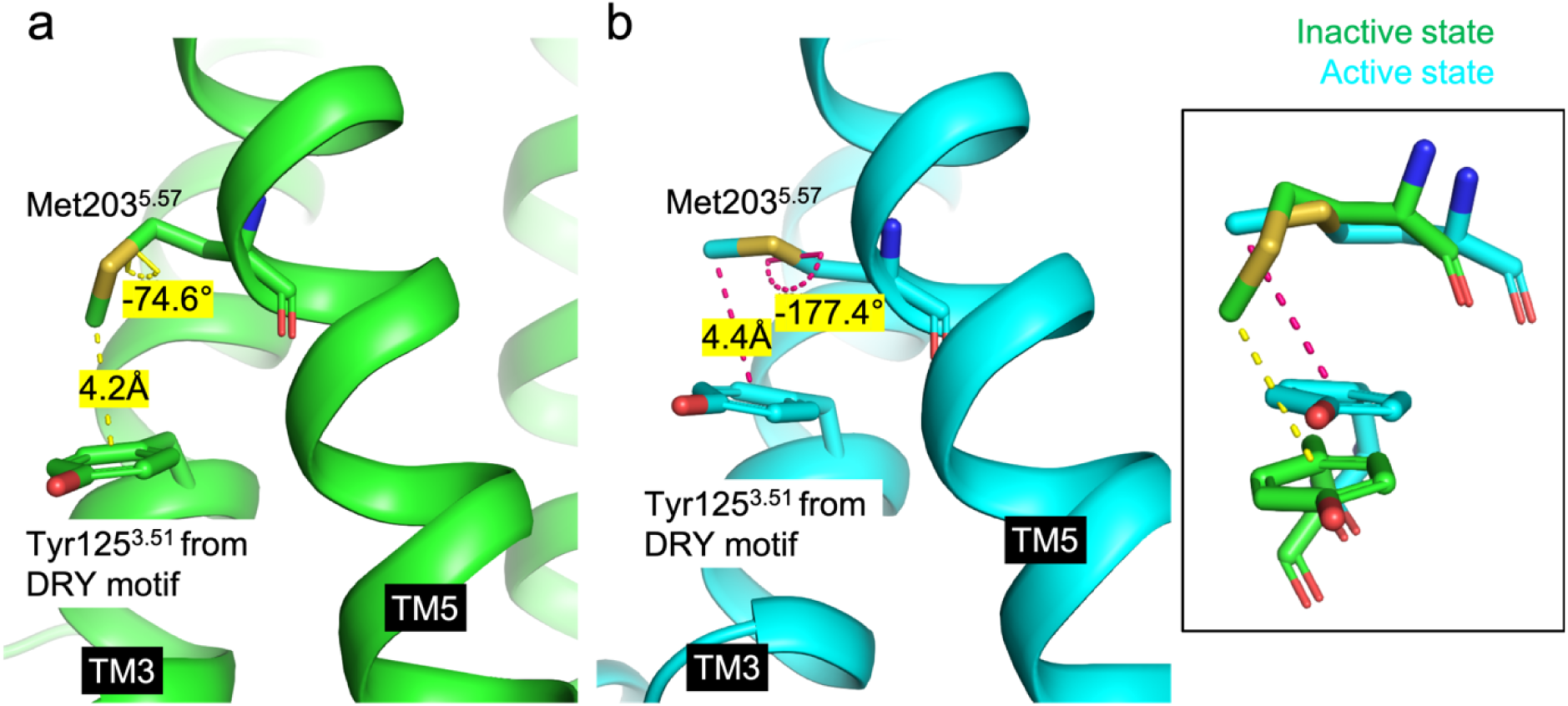
The local environment of Met203^5.57^ and its χ3 dihedral angle. The methyl group of Met203^5.57^ sits on top of Tyr125^3.51^ of the DRY motif as shown in the α_1A_-AR-A4 homology models, and is expected to experience a ring current effect from Tyr125^3.51^. (a) In the inactive state of α_1A_-AR-A4 model (green), the distance between the methyl of Met203^5.57^ and the ring of Tyr125^3.51^ is 4.2Å. The χ3 dihedral angle of Met203^5.57^ is -74.6°, which means the χ3 in the inactive state is averaging between gauche and trans conformers. (b) In the active state of α_1A_-AR-A4 model (cyan), the distance between the methyl of Met203^5.57^ and the ring of Tyr125^3.51^ is 4.4Å. The χ3 dihedral angle of Met203^5.57^ is -177.4°, which means the χ3 in the active state is in a near trans conformer.

**Supplementary Figure 5.**
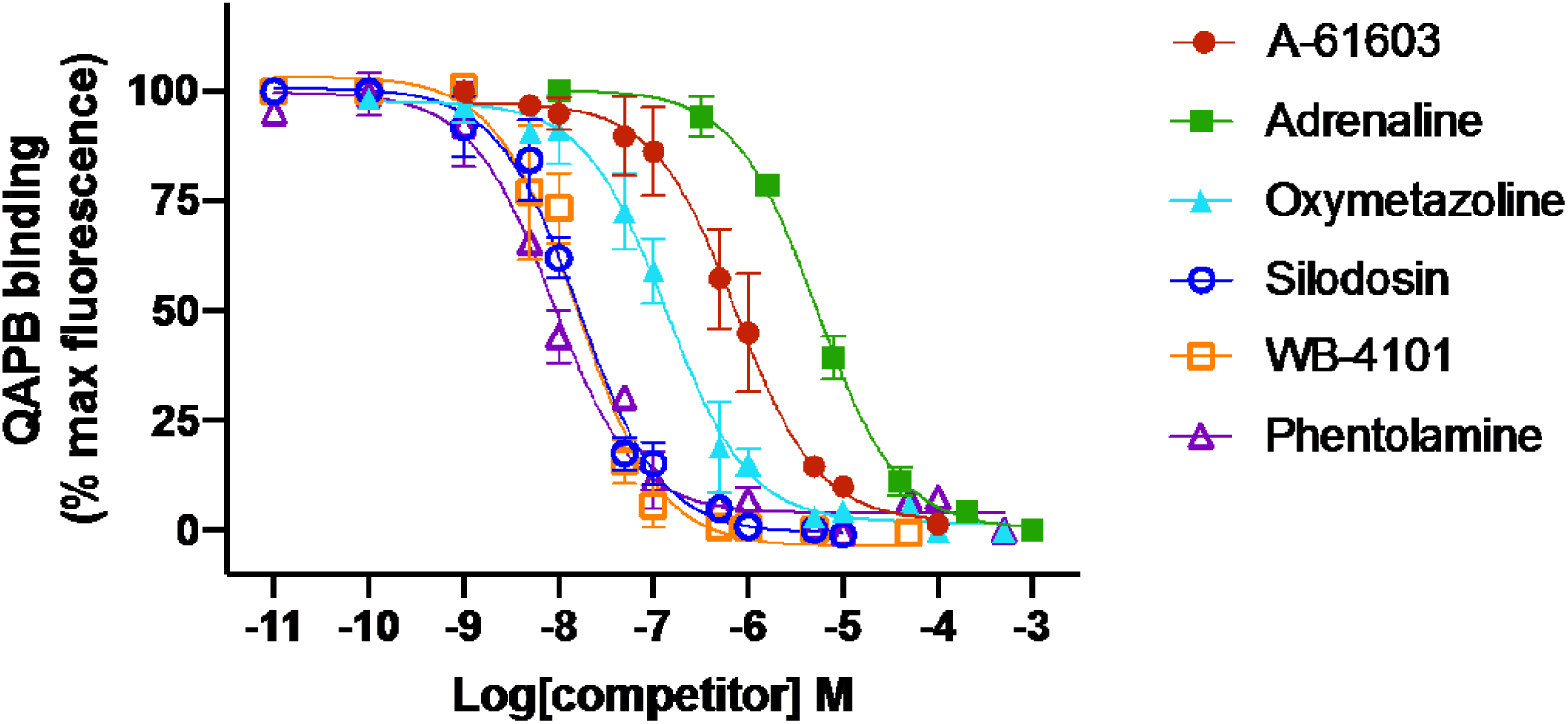
Characterisation of ligand affinity to α_1A_-AR-A4 (L312F). QAPB competition binding for 2 hours at 22 °C against purified α_1A_-AR-A4 (L312F) with A-61603 (maroon, solid circles), adrenaline (green, solid squares), oxymetazoline (cyan, solid triangles), silodosin (blue, open circles), WB-4101 (orange, open squares) and phentolamine (purple, open triangles).

**Supplementary Table 1.** Measured affinities of ligands utilised in the present study.^a^

| | $\alpha_{1A}$-AR WT | $\alpha_{1A}$-AR-A4 | $\alpha_{1A}$-AR-A4 (L312F) | $\alpha_{1A}$-AR-A4-active |
| --- | --- | --- | --- | --- |
| QAPB $K_D$ (nM) | N.D | $11.6 \pm 2.0^1$ | $30.3 \pm 21.2$ | $29.2 \pm 15.1$ |
| | $K_i^b$ | $K_i$ | $K_i$ | $K_i$ |
| Adrenaline | $3.3 \pm 0.4 \mu M^2$ | $>1.0 mM^1$ | $2.7 \pm 0.5 \mu M$ | $0.4 \pm 0.1 mM$ |
| Phenylephrine | $6.2 \pm 1.5 \mu M^2$ | $\sim 0.6 mM^1$ | $41.9 \pm 26.6 \mu M^1$ | $1.6 \pm 0.6 mM$ |
| A-61603 | $\sim 79.4 nM^3$ | $113.5 \pm 47.2 \mu M^1$ | $0.4 \pm 0.2 \mu M$ | $18.4 \pm 18.9 \mu M$ |
| Oxymetazoline | $6.7 \pm 0.9 nM^4$ | $52.8 \pm 8.0 \mu M$ | $65.7 \pm 24.4 nM$ | $9.7 \pm 6.4 \mu M$ |
| PF-3774076 | $\sim 83.0 nM^5$ | N.D | N.D | $19.4 \pm 12.9 \mu M$ |
| Silodosin | $0.036 \pm 0.010 nM^4$ | N.D | $8.4 \pm 1.9 nM$ | N.D |
| Phentolamine | $2.7 \pm 0.1 nM^4$ | N.D | $3.9 \pm 2.0 nM$ | N.D |
| WB-4101 | $0.21 \pm 0.03 nM^4$ | N.D | $7.6 \pm 3.4 nM$ | N.D |
| Prazosin | $0.17 \pm 0.02 nM^4$ | $57.0 \pm 11.8 nM^1$ | $7.5 \pm 3.8 nM^1$ | N.D |

<sup>a</sup> Data are presented as mean $K_i \pm SD$ and mean $K_D \pm SD$ , except for the data cited from the literature which are mean $K_i \pm SEM$ and mean $K_D \pm SEM$ . Three independent biological replicate experiments (n=3) were done for all data. N.D, not determined. <sup>b</sup>These $K_i$ were measured on cells overexpressed with WT human $\alpha_{1A}$ -AR. $K_i$ of ligands on $\alpha_{1A}$ -AR-A4, $\alpha_{1A}$ -AR-A4 (L312F) and $\alpha_{1A}$ -AR-A4-active were determined with purified receptors using Kingfisher binding assay (see methods). For some ligand-receptor pairings, full displacement in competition binding assays was not observed, and thus only approximate $K_i$ values could be estimated (indicated with ~)

**Supplementary Figure 6.**
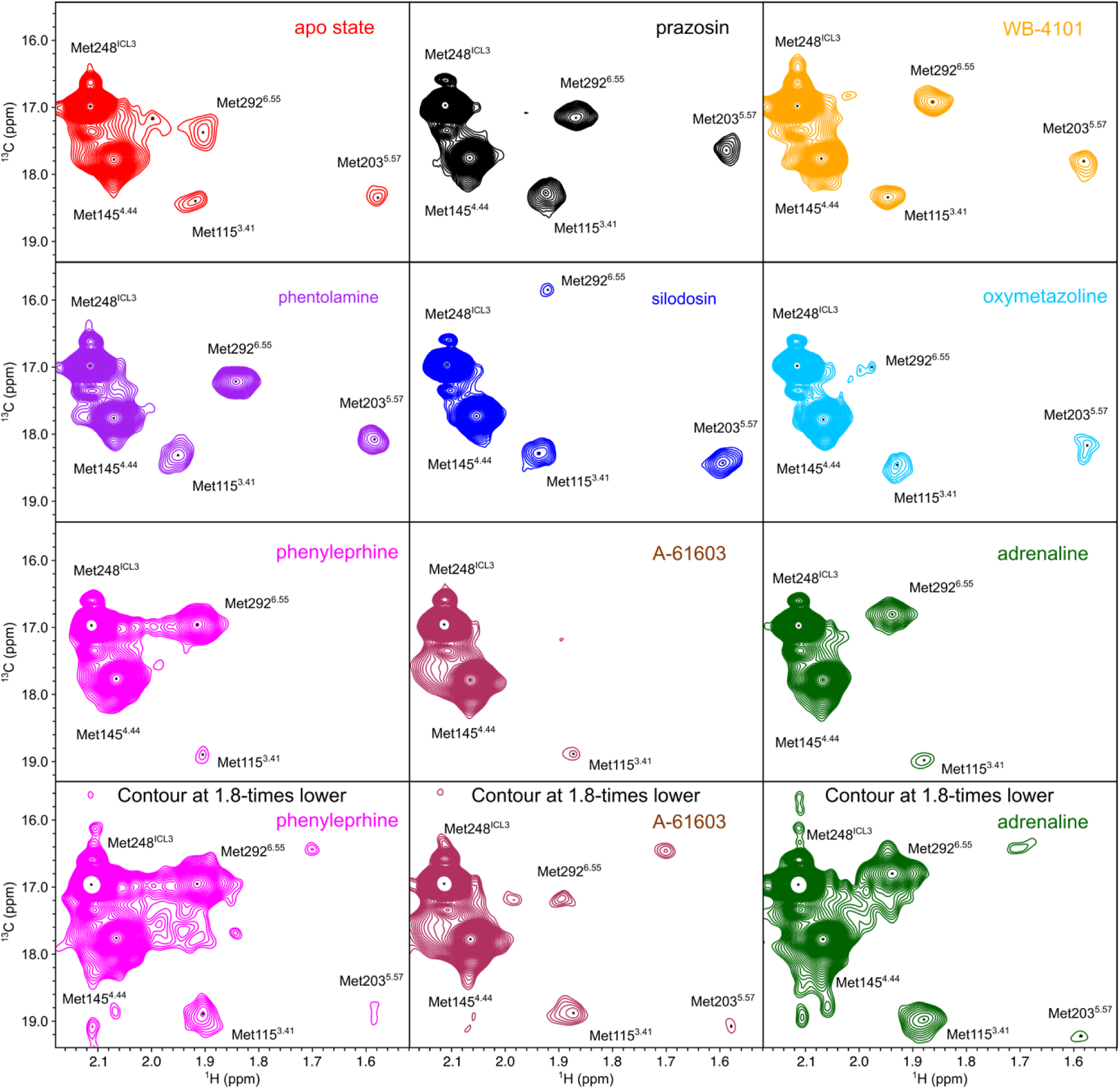
^1^H-^13^C SOFAST-HMQC spectra of α_1A_-AR-A4 (L312F). Individual NMR spectrum for [^13^C^ε^H_3_-Met] α_1A_-AR-A4 (L312F) collected in the apo-state (red) and bound to prazosin (black, inverse agonist), WB-4101 (yellow, inverse agonist), phentolamine (purple, inverse agonist), silodosin (blue, neutral antagonist), oxymetazoline (cyan, partial agonist), phenylephrine (magenta, full agonist), A-61603 (maroon, full agonist), and adrenaline (green, full agonist).

**Supplementary Figure 7.**
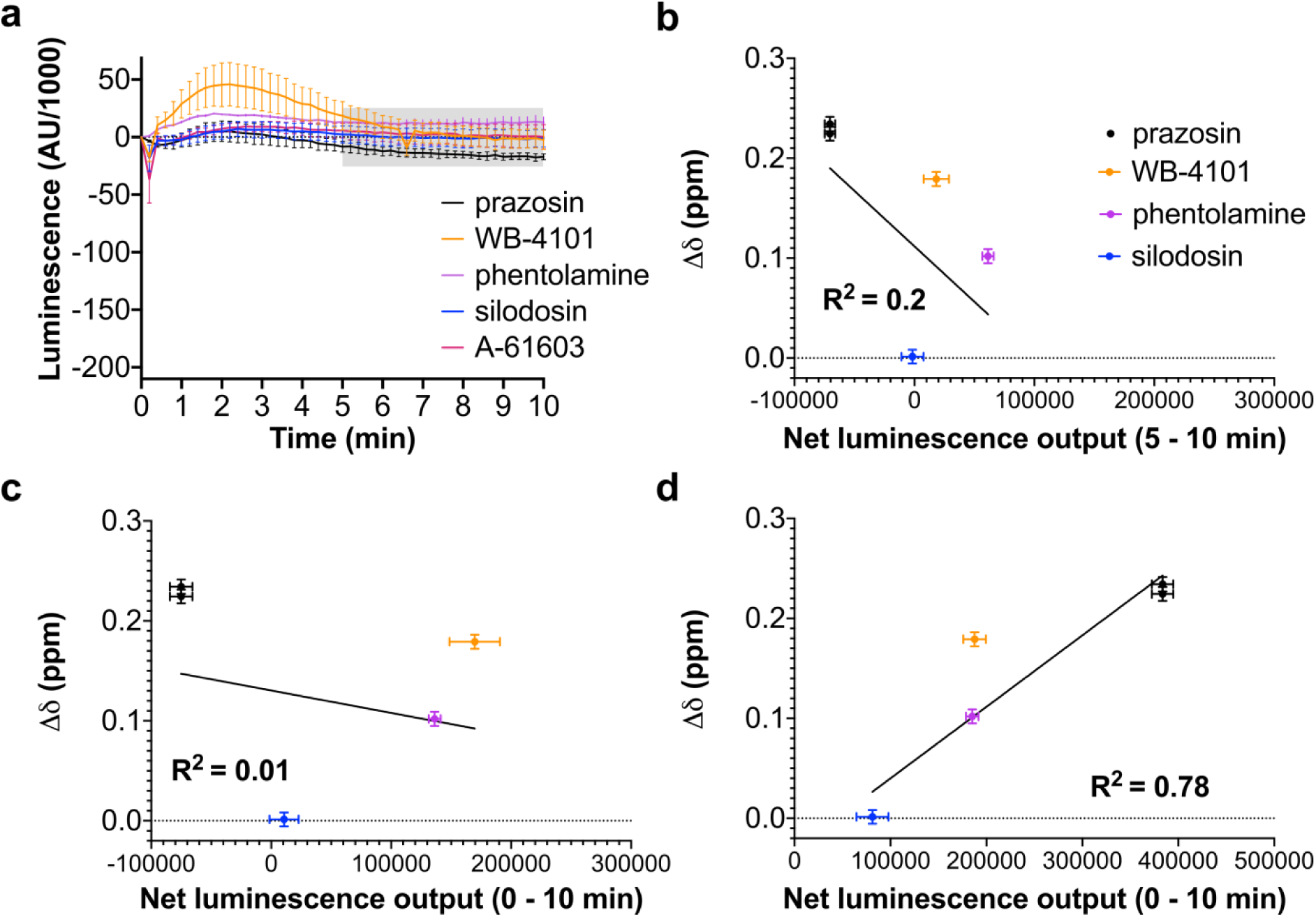
Controls for NanoBit G protein activity assay. (a) NanoBit G protein activity assay on empty vector (pcDNA3.1/Zeo) transfected COS-7 cells treated with the same concentrations of prazosin, WB-4101, phentolamine and silodosin as in Figure 4b. The grey shaded region indicates where the area under the curve measurements were taken for (b). (b) Linear regression analysis of the average chemical shift differences (Δδ) for the ^13^C^ε^H_3_ of Met203 in α_1A_-AR-A4 (L312F) and the increase in luminescence seen between 5 – 10 min after treatment in the NanoBit assay on empty vector (pcDNA3.1/Zeo) transfected COS-7 cells. A P value of 0.2663 was obtained when testing against the null hypothesis of a slope of 0 (c) Linear regression analysis of the average chemical shift differences (Δδ) for the ^13^C^ε^H_3_ of Met203 in α_1A_-AR-A4 (L312F) and the increase in luminescence seen for the first 10 min after treatment in the NanoBit assay on empty vector (pcDNA3.1/Zeo) transfected COS-7 cells. Ligands are coloured as listed above and a P value of 0.6754 indicated slope not deviating significantly from 0. (d) Linear regression analysis of the average chemical shift differences (Δδ) for the ^13^C^ε^H_3_ of Met203 in α_1A_-AR-A4 (L312F) and the increase in luminescence seen for the first 10 min after treatment in the NanoBit assay on COS-7 cells transfected with wild-type α_1A_-AR (as in Figure 4b-c). A P value of 0.0071 suggested a significantly non-zero slope. Ligands are coloured as listed above. In (b), (c) and (d) Δδ are plotted for two independent titrations of prazosin and silodosin, and single experiments for WB-4101 and phentolamine. Average chemical shift differences (Δδ) were normalised using the equation Δδ=[(Δδ_1H_)^2^+(Δδ_13C_/3.5)^2^]^0.5^ and errors were calculated by the formula [Δδ_1H_*R_1H_+Δδ_13C_*R_13C_/(3.5)^2^]/Δδ, where R_1H_ and R_13C_ are the digital resolutions in ppm in the ^1^H and ^13^C dimensions respectively.

**Supplementary Figure 8.**
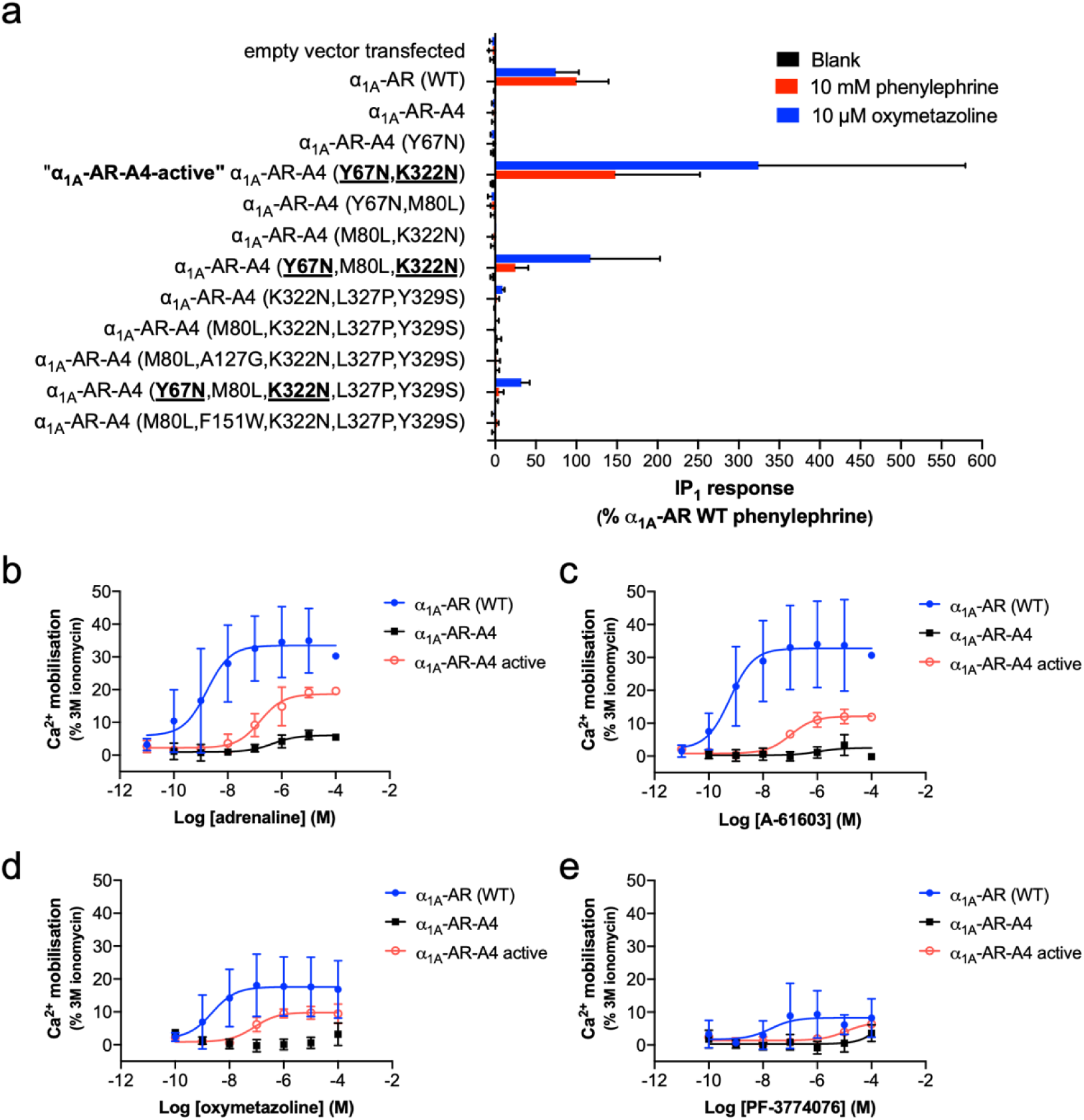
Functional signalling assays performed on α_1A_-AR-A4 active and other mutants. (a) Measurement of agonist (phenylephrine and oxymetazoline) induced accumulation of IP_1_ in COS-7 cells transfected with WT α_1A_-AR, α_1A_-AR-A4 and mutants that were made with reverted mutations on α_1A_-AR-A4. Y67N, M80L, A127G, F151W, K322N, L327P and Y329S are the predicted critical back mutations that were screened to recover the signaling ability of α_1A_-AR-A4. All of α_1A_-AR-A4 back mutants containing Y67N and K322N (highlighted with bold and underlined) displayed accumulation of IP_1_ signal upon agonist activation. α_1A_-AR-A4 (Y67N, K322N) is labelled as α_1A_-AR-A4-active. In this screening assay some mutants were only measured in one biological replicate experiment (α_1A_-AR-A4 (M80L,K322N,L327P,Y329S; α_1A_-AR-A4 (K322N,L327P,Y329S); α1A-AR-A4 (Y67N,M80L,K322N); α_1A_-AR-A4 (M80L,K322N); α_1A_-AR-A4 (Y67N,K322N); α_1A_-AR-A4), with the others measured in two independent biological replicate experiments, with data plotted as mean ± SD of replicate measurements. (b-e) Measurement of adrenaline (b), A-61603 (c), oxymetazoline (d) and PF-3774076 (e) induced Ca^2+^ mobilization in COS-7 cells transfected with α_1A_-AR (blue, solid circles), α_1A_-AR-A4 (black, solid squares) and α_1A_-AR-A4 active (red, open circles). Data represent the mean ± SD from three independent biological replicate experiments, each measured as three technical replicates.

**Supplementary Figure 9.**
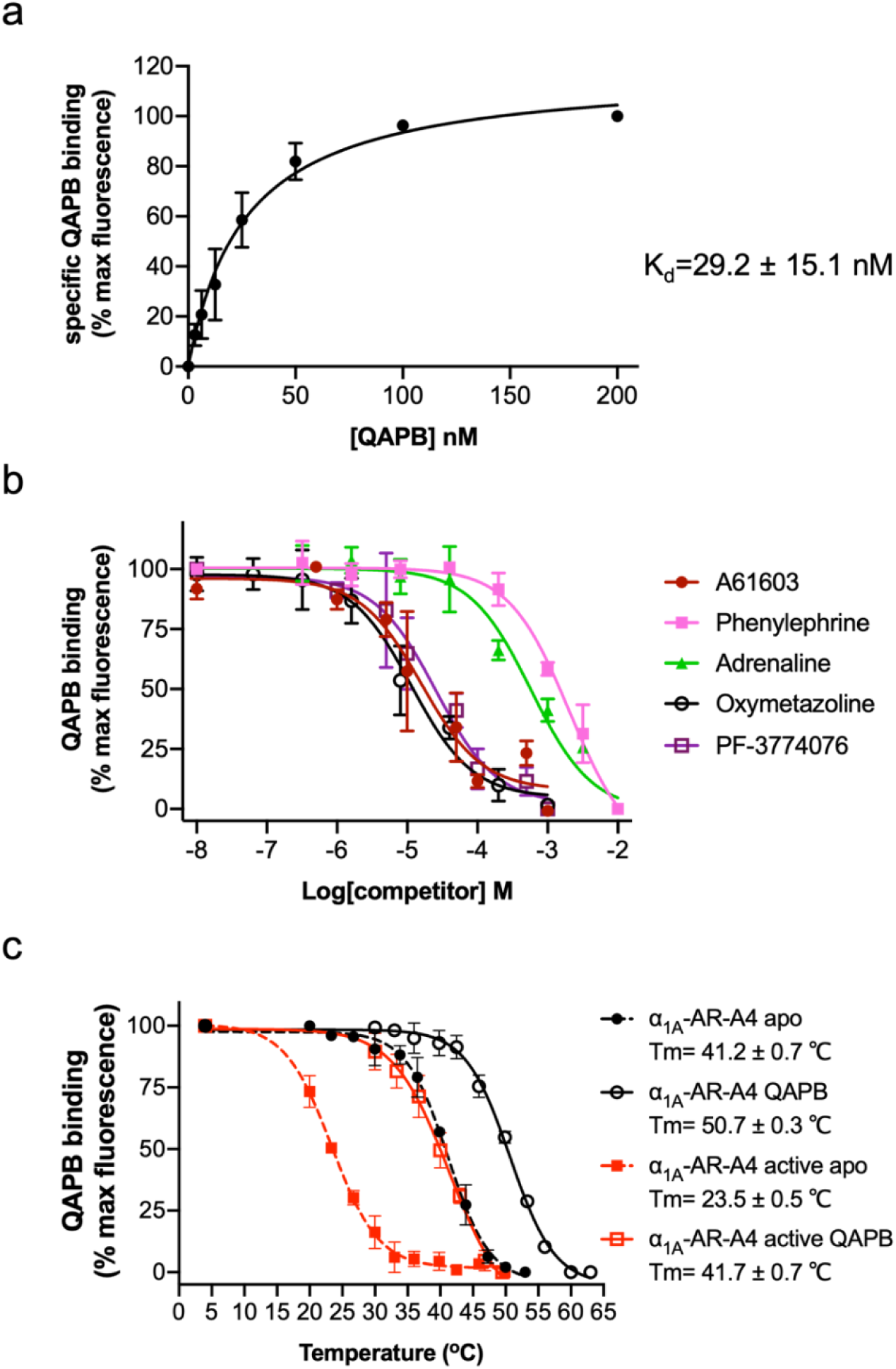
Characterisation of α_1A_-AR-A4-active. (a) Saturation binding of QAPB to purified α_1A_-AR-A4 active. (b) QAPB competition binding for 2 hours at 22 °C against purified α_1A_-AR-A4 active with A-61603 (maroon, solid circles), phenylephrine (pink, solid squares), adrenaline (green, solid triangles), oxymetazoline (black, open circles), PF-3774076 (purple, open squares). (c) Thermostability assay performed on α_1A_-AR-A4 in the apo state (black solid circles and dash line), QAPB-bound state (black open circles and solid line) and α_1A_-AR-A4-active in the apo state (red solid squares and dash line), QAPB-bound state (red open squares and solid line).

**Supplementary Figure 10.**
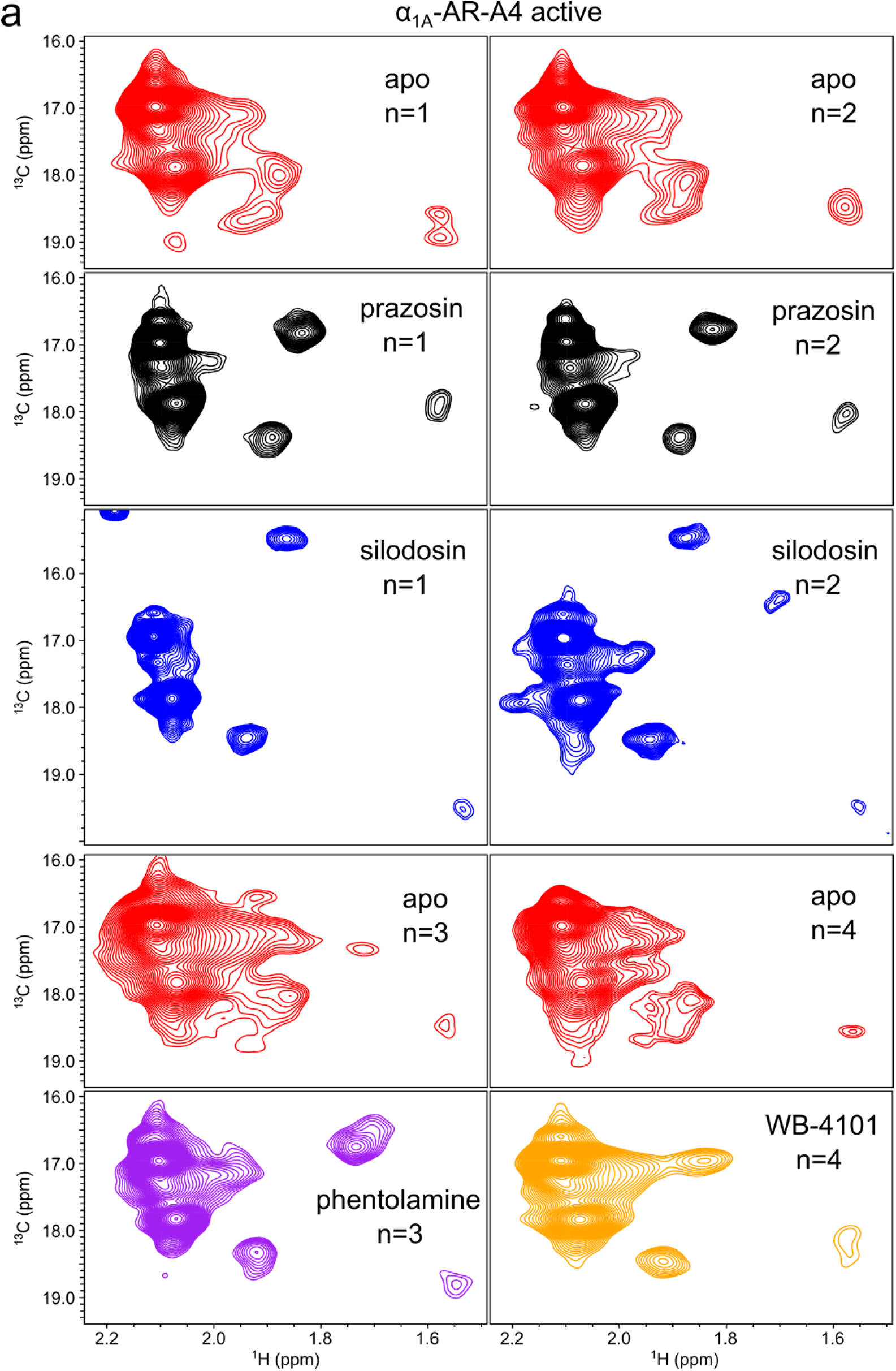
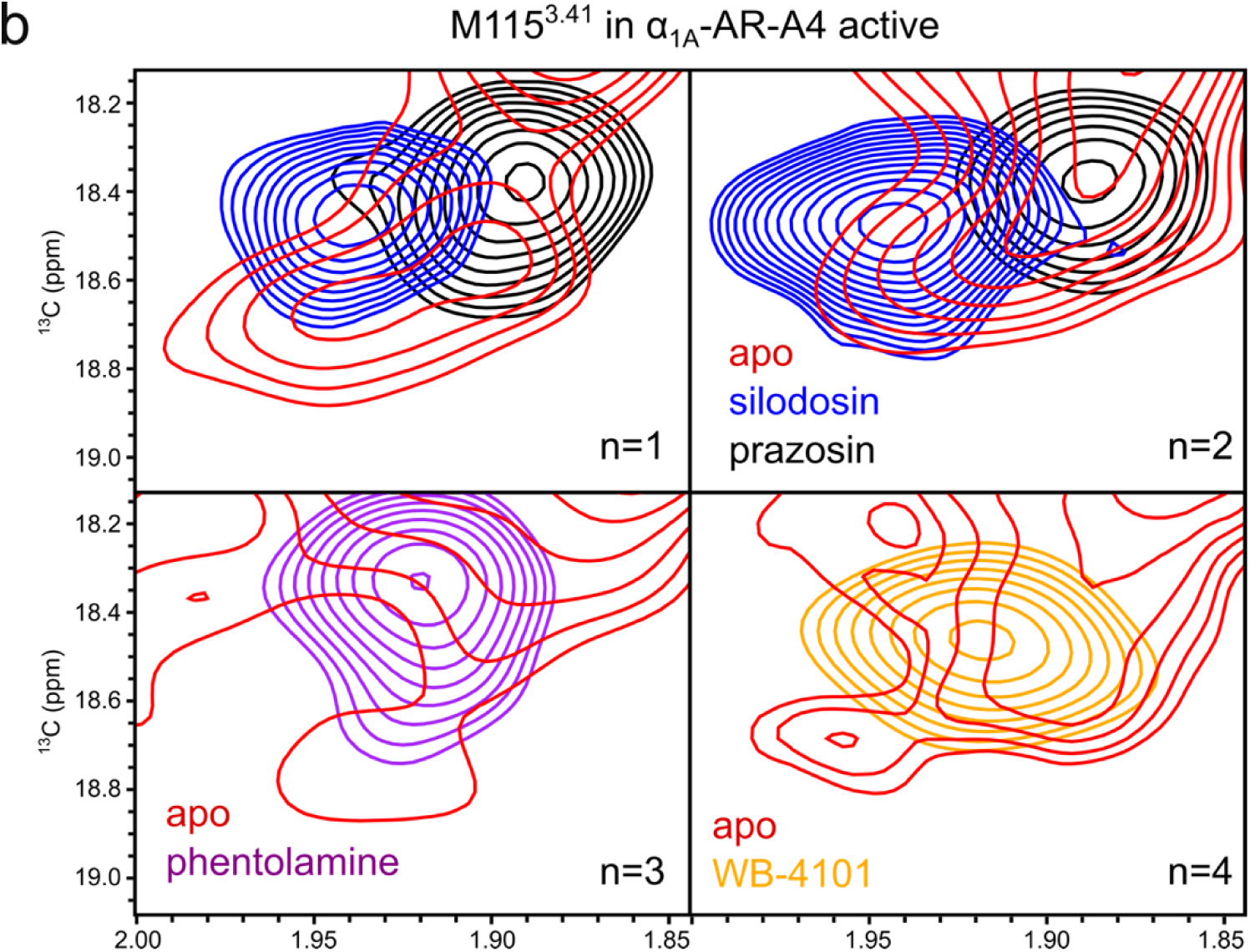
^1^H-^13^C SOFAST-HMQC spectra of α_1A_-AR-A4-active. (a) Four separate expressions and purifications of α_1A_-AR-A4-active were conducted and data acquired for apo-(red), prazosin (black) and silodosin (blue), phentolamine (purple) and WB-4101 (orange). (b) Expansions and overlay of the region where the ^13^C^ε^H_3_ of Met115 resonates.

**Supplementary Figure 11.**
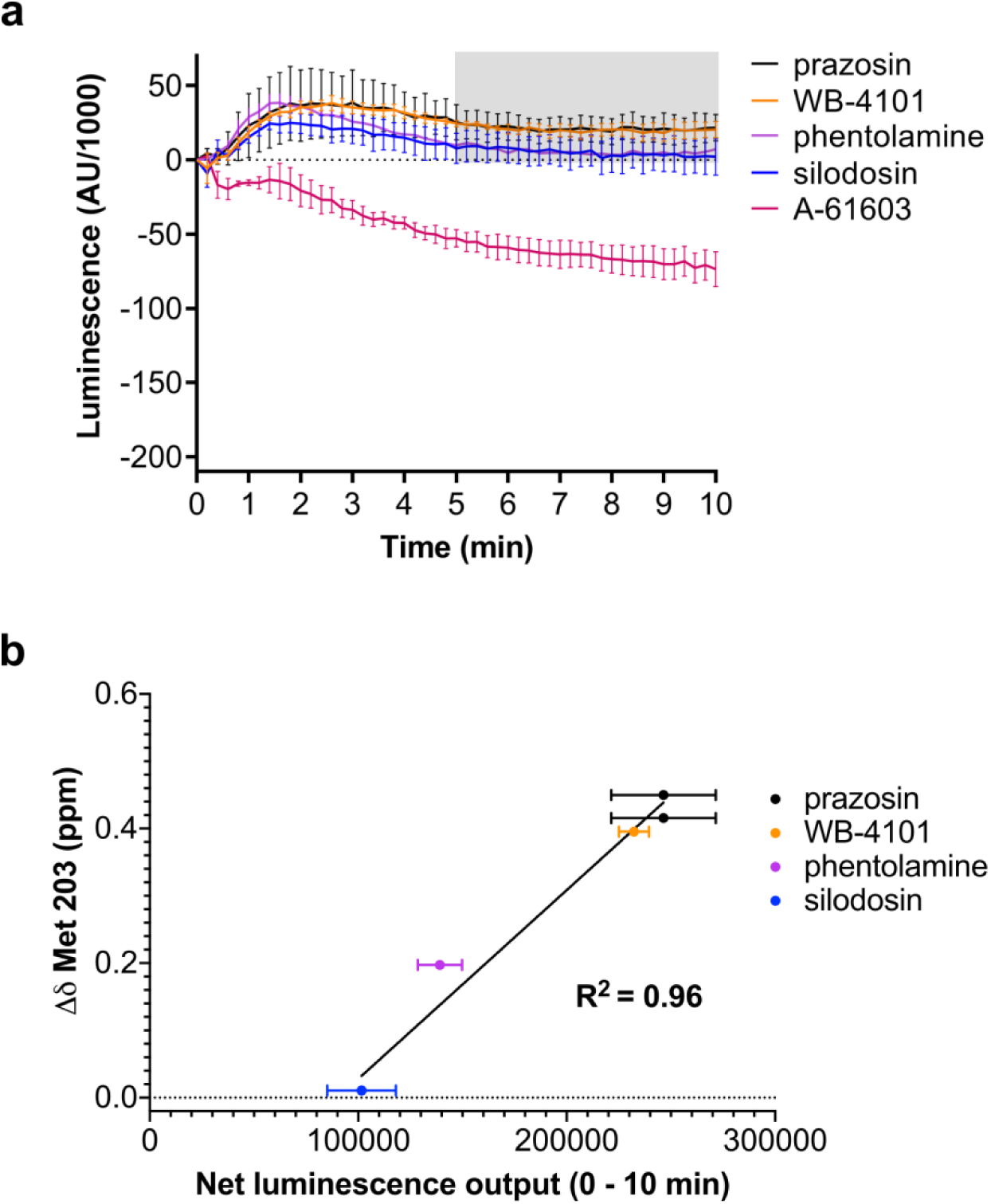
Correlation between the chemical shift position of Met203^5.57^ in α_1A_-AR-A4-active and inverse agonists efficacy. (a) NanoBit G protein activity assay demonstrating inverse agonism of prazosin, WB-4101, phentolamine and silodosin at α_1A_-AR-A4 active expressing COS-7 cells. The grey shaded region indicates where the area under the curve measurements were taken to make Figure 5e. (b) Linear regression analysis of the average chemical shift differences (Δδ) for the ^13^C^ε^H_3_ of Met203 in α_1A_-AR-A4-active and the increase in luminescence seen over the first 10 minutes in the NanoBit assay for each antagonist. A P value of 0.0002 suggested that the slope was significantly different from zero. In (b) Δδ are plotted for two independent titrations of prazosin and silodosin, and single experiments for WB-4101 and phentolamine. Average chemical shift differences (Δδ Met 203) were normalised using the equation Δδ=[(Δδ_1H_)^2^+(Δδ_13C_/3.5)^2^]^0.5^ and errors were calculated by the formula [Δδ_1H_*R_1H_+Δδ_13C_*R_13C_/(3.5)^2^]/Δδ, where R_1H_ and R_13C_ are the digital resolutions in ppm in the ^1^H and ^13^C dimensions respectively.

